# Resolving the *Euphorbia neriifolia* (Euphorbiaceae) complex: a taxonomy with two new succulent species from Western India

**DOI:** 10.1101/2024.04.18.590052

**Authors:** Nilesh V. Malpure, Bruce E. De Jong, Prashant S. Raut, Milind M. Sardesai

## Abstract

Linnaeus appears to have clubbed three distinct species under *Euphorbia neriifolia*, a species from China and Indonesia that is the original *E. neriifolia*, a species from India and Sri Lanka which is now recognized as *E. nivulia*, and a species from West Africa which is now called *E. drupifera*. After conducting field studies and examining the applicable protologues, original type material, world literature, and world herbaria, we conclude that for hundreds of years, workers have erroneously clubbed various species from India under *E. neriifolia*. These species include *E. ligularia* from Northeast India and two common succulents from Western India described here as new species viz. *E. yadavii* and *E. paschimia*. An epitype for *E. neriifolia* from Indonesia is designated to stabilize the species, *E. ligularia* is reinstated as a species, a neotype is designated for *E. drupifera*. *E. complanata* is synonymized under *E. neriifolia*, and *E. undulatifolia* is synonymized under *E. ligularia*. We present amended descriptions of *E. neriifolia*, *E. nivulia*, *E. drupifera* and *E. ligularia*.

## Introduction

The genus *Euphorbia* Linnaeus (1753: 450) has approximately 2000 accepted species (Dorsey *et al*. 2013). Based on a combination of key botanical features and molecular phylogenetic studies, genus *Euphorbia* was divided into 4 sub-genera of which sub-genus *Euphorbia* has 661 species (Dorsey *et al*. 2013). This sub-genus is further divided into 4 clades and 21 sections. Of these sections, *Euphorbia* sect. *Euphorbia* consists of approximately 339 species, most of which possess the characteristic spine shield and all of which are succulents.

In India, Binojkumar & Balakrishnan (2010) reported 84 species in genus *Euphorbia.* Under *E.* section *Euphorbia* there are currently 22 Indian species by our count; the 11 species recognised by Binojkumar & Balakrishnan (2010) (excluding *E. lactea* Haworth (1812:127) and *E. trigona* Miller (1768: No.3) which are non-Indian), the 4 recognized Indian geophytes (Mane *et al*. 2017), *E. epiphylloides* Kurz (1873: 247) from the Andamans, and several recently described taxa viz. *E. gokakensis* Malpure *et al*. (2016: 380), *E. venkatarajui* Sarojinidevi (2017: 359), *E. belagaviensis* Sarojinidevi & Kullayiswamy (2018: 24), *E. lakshminarasimhanii* Malpure et. al. (2021a: e03142), *E. sahyadrica* Malpure *et. al.* (2021b: 285), *E. ravii* Swamy & Prasad (2022: 229).

Our recent field studies in India and Indonesia have demonstrated that the Indian plants widely considered to be *E. neriifolia* (Binojkumar & Balakrishnan 2010) are very different from the Indonesian plants, also widely considered to be *E. neriifolia*. Moreover, an extensive examination of the literature shows that there has been, and continues to be, considerable confusion as to what exactly constitutes this species. Because the name *E. neriifolia* has often been erroneously and imprecisely applied for over 250 years, it is vital to establish the type characteristics of the species and understand what plant material Linnaeus was working with when he first described this taxon. On the basis of this, we can then analyse the other members of this complex.

### Historical analysis of the *E. neriifolia* complex

In the protologue of *Euphorbia neriifolia,* Linnaeus (1753) states the following:

1. “Euphorbia aculeata seminuda: angulis oblique tuberculatis” [Spiny euphorbia, partially without leaves [having] oblique, tuberculate angles], referencing Linnaeus (1737: 196, 1747: 90, 1748: 139), van Royen (1740: 194), and Wiman (1752: 109, no. 7).
2. “Euphorbium afrum spinosum, foliis latioribus non spinosis” [Spiny African euphorbia [with] broad spineless leaves], referencing Seba (1734: 18, plate 9, fig. 1). [Fig.1 A]
3. “Tithymalus aizoides arborescens, spinosus, caudice angulari, nerii folio” [Evergreen spurge becoming treelike [with a] spiny, angled trunk, leaves like oleander], referencing C. Commelin (1703: 56, fig. 6). [Fig. 1 B]
4. “Tithymalus indicus spinosus, nerii folio” [Spiny Indian spurge [with] leaves like oleander], referencing J. Commelin (1697: 25, plate 13). [Fig. 1 C]
5. “Ela calli” [Leafy spurge, translated from Malayalam and Tamil], referencing (Rheede 1689: 83, plate 43). [Fig. 1 D]
6. “Habitat in India”.

**FIGURE 1.**
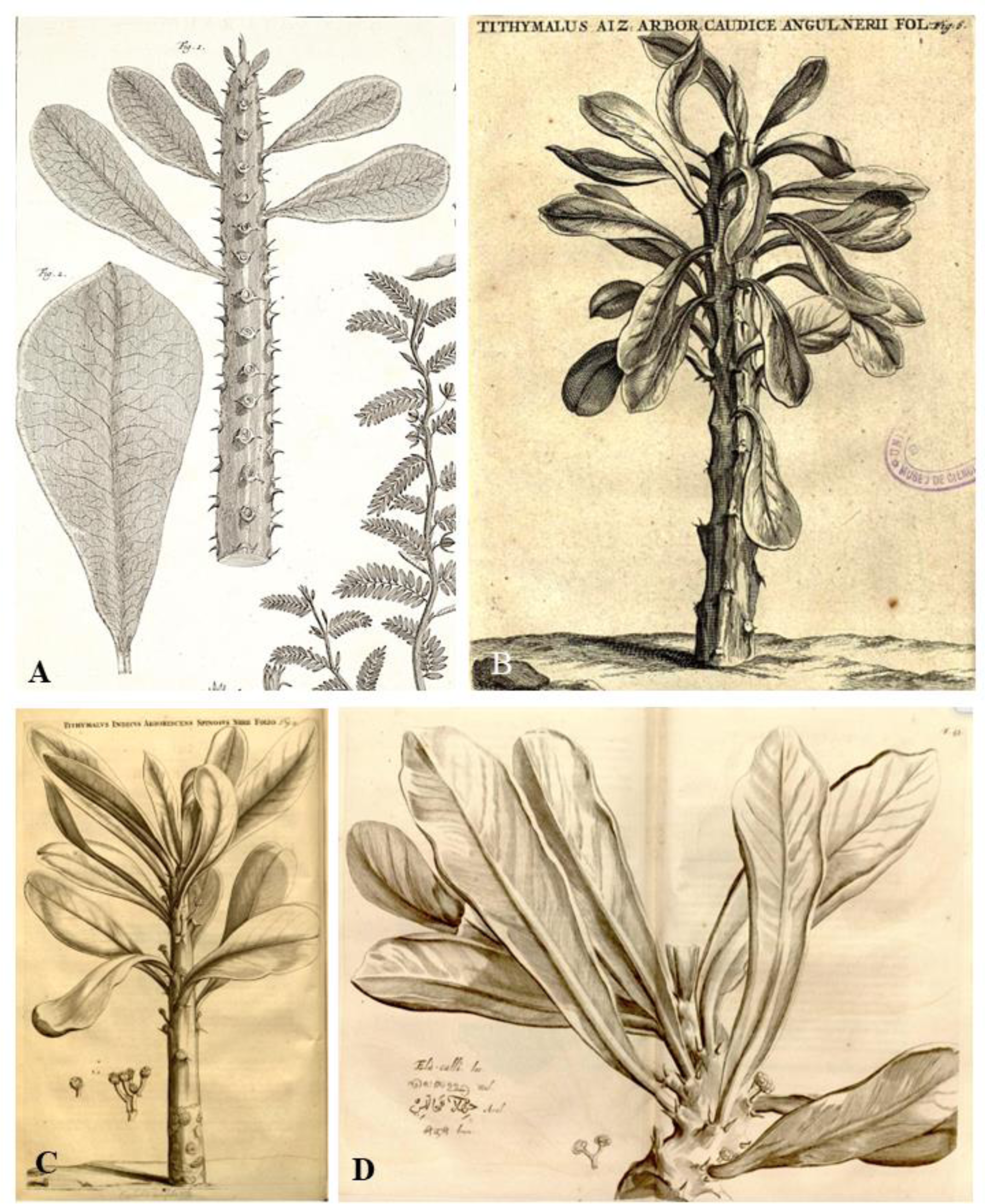
Fig. 1 from Table IX, Seba (1734). In public domain. Fig. 6, C. Commelin (1703). Used with permission. Digital Library of the Royal Botanic Garden, RJB-CSIC: https://bibdigital.rjb.csic.es Plate 13, J. Commelin (1697). In public domain. Plate 43, Rheede tot Draakestein (1679). In public domain.

**TABLE 1.**
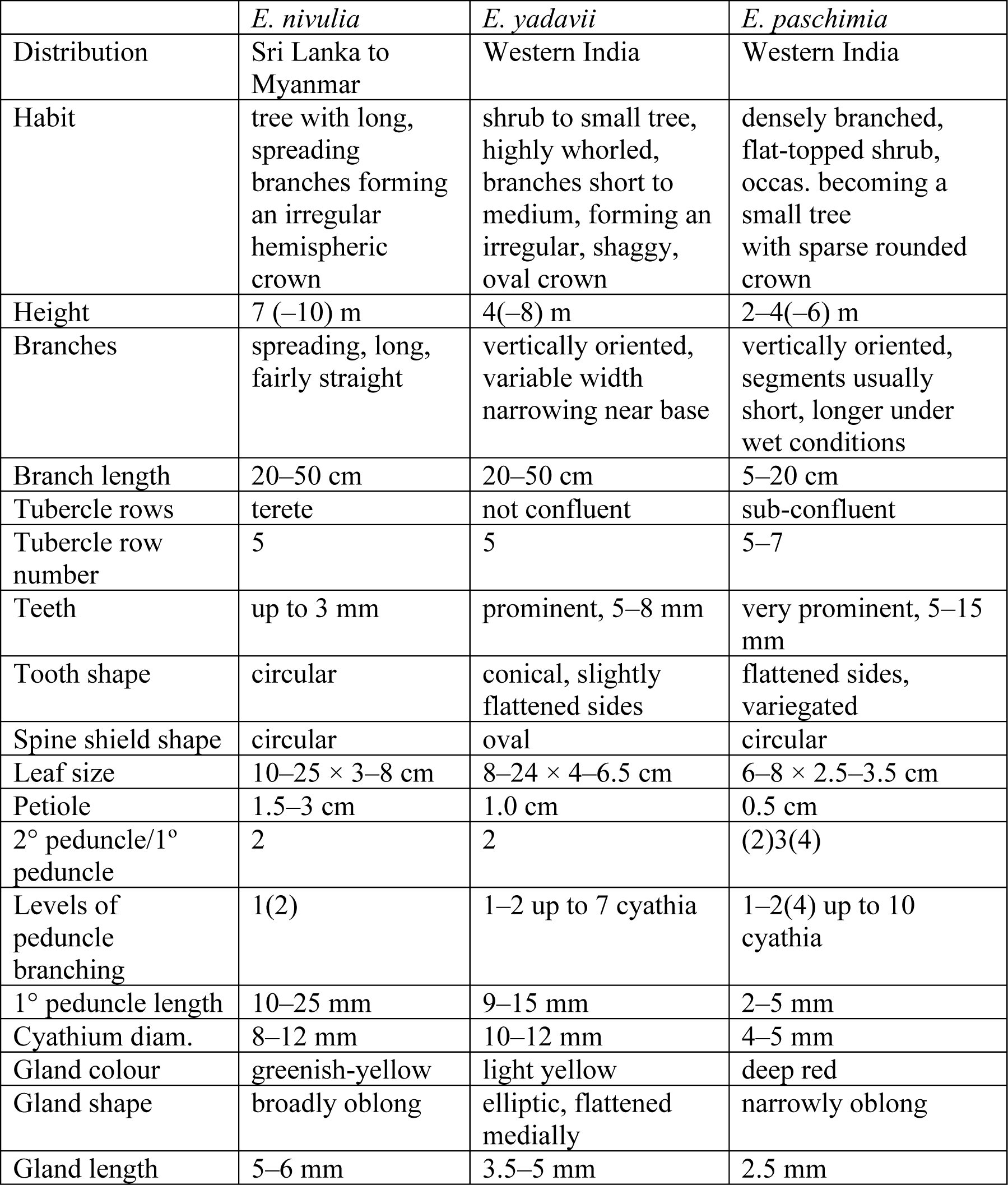
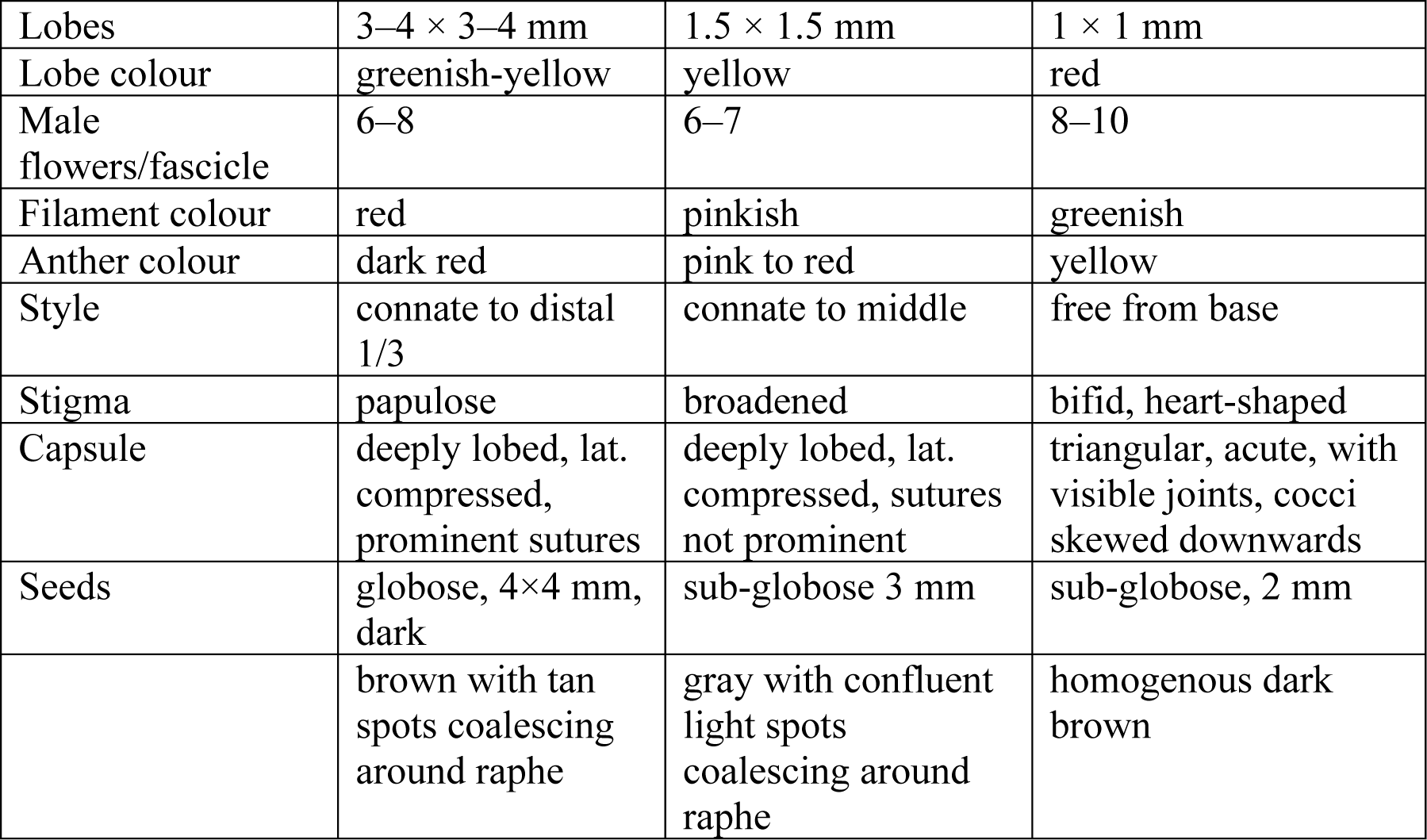
Comparison of Euphorbia nivulia, Euphorbia yadavii, and Euphorbia paschimia.

|  | *E. nivulia* | *E. yadavii* | *E. paschimia* |
| --- | --- | --- | --- |
| Distribution | Sri Lanka to Myanmar | Western India | Western India |
| Habit | tree with long, spreading branches forming an irregular hemispheric crown | shrub to small tree, highly whorled, branches short to medium, forming an irregular, shaggy, oval crown | densely branched, flat-topped shrub, occas. becoming a small tree with sparse rounded crown |
| Height | 7 (–10) m | 4(–8) m | 2–4(–6) m |
| Branches | spreading, long, fairly straight | vertically oriented, variable width narrowing near base | vertically oriented, segments usually short, longer under wet conditions |
| Branch length | 20–50 cm | 20–50 cm | 5–20 cm |
| Tubercle rows | terete | not confluent | sub-confluent |
| Tubercle row number | 5 | 5 | 5–7 |
| Teeth | up to 3 mm | prominent, 5–8 mm | very prominent, 5–15 mm |
| Tooth shape | circular | conical, slightly flattened sides | flattened sides, variegated |
| Spine shield shape | circular | oval | circular |
| Leaf size | 10–25 × 3–8 cm | 8–24 × 4–6.5 cm | 6–8 × 2.5–3.5 cm |
| Petiole | 1.5–3 cm | 1.0 cm | 0.5 cm |
| 2° peduncle/1° peduncle | 2 | 2 | (2)3(4) |
| Levels of peduncle branching | 1(2) | 1–2 up to 7 cyathia | 1–2(4) up to 10 cyathia |
| 1° peduncle length | 10–25 mm | 9–15 mm | 2–5 mm |
| Cyathium diam. | 8–12 mm | 10–12 mm | 4–5 mm |
| Gland colour | greenish-yellow | light yellow | deep red |
| Gland shape | broadly oblong | elliptic, flattened medially | narrowly oblong |
| Gland length | 5–6 mm | 3.5–5 mm | 2.5 mm |
| Lobes | 3–4 × 3–4 mm | 1.5 × 1.5 mm | 1 × 1 mm |
| Lobe colour | greenish-yellow | yellow | red |
| Male flowers/fascicle | 6–8 | 6–7 | 8–10 |
| Filament colour | red | pinkish | greenish |
| Anther colour | dark red | pink to red | yellow |
| Style | connate to distal 1/3 | connate to middle | free from base |
| Stigma | papulose | broadened | bifid, heart-shaped |
| Capsule | deeply lobed, lat. compressed, prominent sutures | deeply lobed, lat. compressed, sutures not prominent | triangular, acute, with visible joints, cocci skewed downwards |
| Seeds | globose, 4×4 mm, dark | sub-globose 3 mm | sub-globose, 2 mm |
|  | brown with tan spots coalescing around raphe | gray with confluent light spots coalescing around raphe | homogenous dark brown |

Linnaeus did not cite any type material in the protologue. However, he did prepare a sheet labeled 630.1 (The Linnaean Collections, N.D.), annotated in his hand with “7” (in reference to this being the 7^th^ plant under genus *Euphorbia*), “nerifolia”, “Chin(e)”, and on the reverse “Chinensibus inseruit pro sepibus” [Grown by the Chinese as hedges]. This was designated as the lectotype of *E. neriifolia* L. by Radcliffe-Smith (1982b: 93) [Fig. 2 A]. Wijnands (1983) had designated ‘fig. 6 by C. Commelin’ as the lectotype, but this is superseded by the earlier designation of Radcliffe-Smith.

**FIGURE 2.**
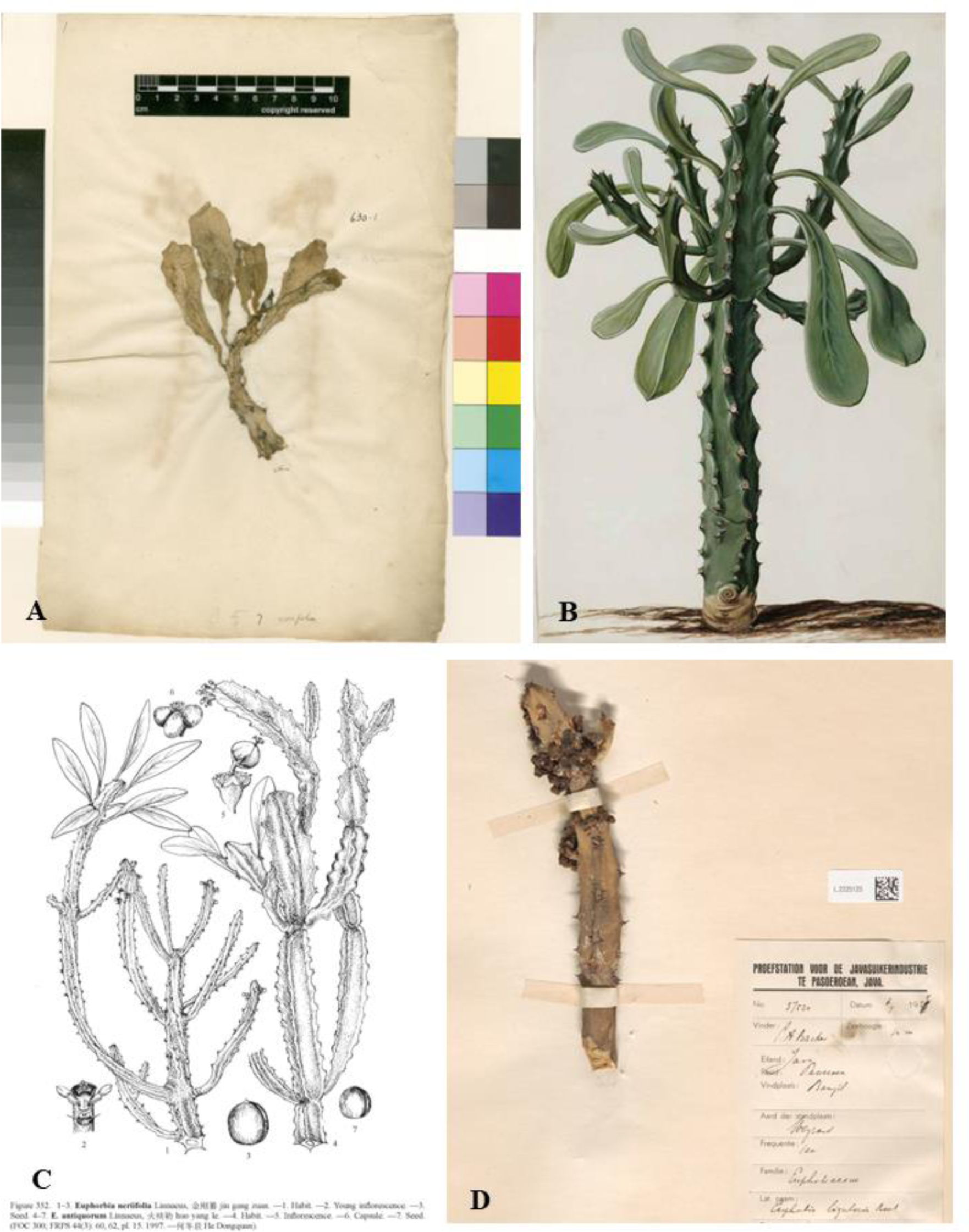
LINN 630.1, Used with permission. The Linnean Society of London. Plate 45, Vol. 5 of the Moninckx Atlas, J. Moninckx (1690). Used with permission. University of Amsterdam, Allard Pierson Depot, OTM: hs. VI G 5 Fig. 352, Flora of China Vol. 11, p. 352. Used with the permission of the Missouri Botanical Garden Press, St. Louis and the Science Press, Beijing. L2225125. Used with permission of Senior Collections Manager, Naturalis Biodiversity Center.

Though not stated explicitly, the notations on the herbarium sheet LINN 630.1 and the correspondence between Linnaeus and Osbeck indicates that this specimen was most likely collected by Osbeck in China in Guangdong “Canton” [Guangzhou] on 10 Sept 1751 (Osbeck 1757; 1771; Savage 1945; Hansen & Maule 1973; Jarvis 2007), and personal communication with the Head of Collections and Curator (Plants) Linnaean Herbarium.

Furthermore, the literature reveals that the Seba plant [Fig. 1 A] had been collected in West Africa and brought to Europe. These plants had been growing in Beaumont’s garden (Croizat 1934, Wijnands 1987), were described there by Kiggelaer (1690), and illustrated in the same place by Seba (1734). This plant was identified as *Elaeophorbia drupifera* Schum. & Thonn. (Croizat 1934: 42), a species now accepted as *Euphorbia drupifera* Thonning (1827: 250).

The plant referenced as C. Commelin (1703) [Fig. 1 B] had been given to Commelin in Hortus Medicus Amsterdam by Agnes Block who had received it from Ambon Island, Indonesia (C. Commelin 1703: 56). Another illustration of this plant from the same time and place is by Jan Moninckx in 1700–1702 (Wijnands 1983, J. Moninckx 1690). [Fig. 2 B]. Our study of living and herbarium specimens from S.E. Asia and China (see additional specimens examined under *E. neriifolia* below), and the Flora of China illustration from Ma & Gilbert (2008:300, Fig 352) [Fig. 2 C] suggest that these plants are identical to the accepted lectotype from China.

In reference to the J. Commelin (1697) plant [Fig 1 C], J. Commelin, the commissioner of Hortus Medicus in Amsterdam, states that the specimen had been received from the Governor of Colombo [Sri Lanka], Laurentius Pijl. Paul Hermann, a German physician and botanist from the same era, described a similar plant during his botanical collecting in Sri Lanka from 1672-1677 (Herman 1717: 54). The features of these plants correspond to what was separated out from *E. neriifolia* as *E. nivulia* Buchanan-Hamilton (1825: 286).

The fourth *Euphorbia* specimen cited by Linnaeus was “*Ela calli*” (Rheede 1679) [Fig. 1 D]. Hortus Malabaricus was a masterpiece produced by an Indo-European team that catalogued the medicinal plants of Malabar, modern day Kerala, India. It is not known whether living specimens of *Ela-calli* from this collection reached European gardens, but this plant is identical to the J. Commelin plant, i.e. *E. nivulia*.

A year after publishing *Species Plantarum*, Linnaeus (1754) cited the *E. ligularia* of Rumphius (1743: 91, Tab 40) from Indonesia as a pre-Linnaean synonym of *E. neriifolia*. This plant [Fig. 3 A] is identical to the C. Commelin (1703) plant [Fig. 1 B]. After correlating each of these Linnaean references with their original material, geography, associated herbarium collections, and literature, it seems clear that Linnaeus worked with only two Asian species while describing *E. neriifolia*: the plants from China and Indonesia and *E. nivulia* from Sri Lanka and India. In our field work, we have identified three other Indian members of this complex which are completely different from *E. neriifolia* found in China and Indonesia and from *E. drupifera*. The first is *E. ligularia* Roxb. ex Buchanan-Hamilton (1825: 285) from Northeast India which has widely been considered a synonym of *E. neriifolia* but is, in fact, quite distinct. The second and third are fairly common succulents from Western India which we describe here as *E. paschimia* Malpure, Sardesai & B. DeJong, *sp. nov.* and *E. yadavii* Malpure, Sardesai & B. DeJong, *sp. nov*.

**FIGURE 3.**
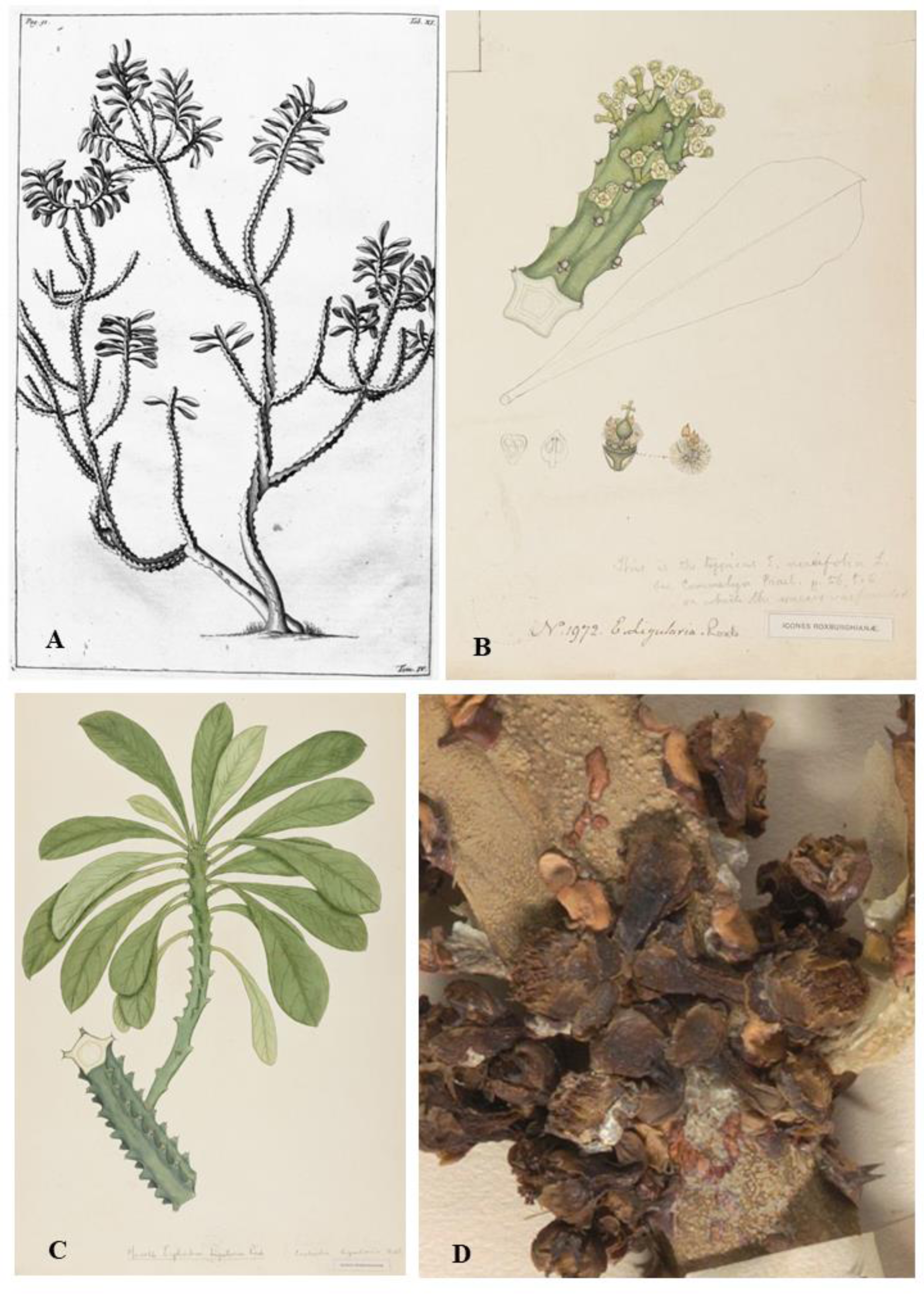
Fig. 11, Vol. 4, Rumphius (1743). Used with permission. Digital Library of the Royal Botanic Garden, RJB-CSIC: https://bibdigital.rjb.csic.es No. 1972, *Euphorbia ligularia* Roxb. from Icones Roxburghianae. Used with permission. © Copyright the Board of Trustees of the Royal Botanical Gardens, Kew. No. 1066, *Euphorbia ligularia* Roxb. from Icones Roxburghianae. Used with permission. © Copyright the Board of Trustees of the Royal Botanical Gardens, Kew. L2225125, close-up view of cyathia. Used with permission of Senior Collections Manager, Naturalis Biodiversity Center.

## Material and Methods

We examined the relevant literature concerning the *E. neriifolia* complex (see below). We also examined the herbaria collections of BAMU, BSI, CAL, DD, JCB, MH, RHT, PDA and SUK. In addition, we studied collections available online from the herbaria ANDA, B, BR, F, M, MICH, L, LINN, NY, TAMA, U, and WAG. Herbarium specimens were assigned to species groupings based on morphology, rather than given labels, since these labels often conflict with morphology.

In Indonesia, we studied plants at Bayview Gardens, Labuan Bajo, Flores as well as populations growing in Pemuteran, Bali below the temple Pura Bato Kursi and in fence lines near Amed Beach, Bali.

In Assam we studied a tree in a private home (300 m elevation close to Kaziranga Park Central Gate, Kohora), an old tree at Iora Resort (about 2 km east of Kaziranga Park Central Gate, Kohora), and a number of home shrines with trees in Bodo Bazaar, Chirang District. For our description, we followed Roxburgh (1832) supplemented by our fresh observations.

Plants from Cote d’Ivoire were studied in various nurseries and home gardens in Abidjan. For the characteristics, we combined observations from these plants, and followed the protologue (Thonning 1827), as well as Staph (1909), Carter (2002), and Schmelzer (2008).

In Tamil Nadu (India), we selected *E. nivulia* samples from the Yercaud and Kalrayan Hills of Tamil Nadu as the basis against which to compare the other species. Specimens included herbarium collections, and plants collected from the above hill ranges. We amended Binojkumar & Balakrishnan’s (2010) descriptions with our own field observations.

In Maharashtra (India), we collected plants from Chandanpuri Ghat, near Sangamner, District Ahmednagar, described here as *E. paschimia*. In addition, we studied plants collected from a fence line in Rampurwadi near Kopargaon, District Ahmednagar and other populations in the region, described as *E. yadavii*.

**17^th^ and 18^th^ Century Literature Examined (by year):—** Rheede (1679), Ray (1688), Breyn (1689), Kiggelaer (1690), M. Moninckx (1690), Plukenet (1696), J. Commelin (1697), Hermann (1698), Morison (1699), J. Moninckx (1702), C. Commelin, (1703), Hermann (1717), Isnard (1720), Tilli (1723), Bradley (1725), Boerhaave (1727), Seba (1734), Aubriet (1690 – 1735), J. Burman (1737), Linnaeus (1737), van Royen (1740), Rumphius (1743), Wiman (1752), Linnaeus (1747, 1748,1753,1754), Osbeck (1757, 1771), Miller (1763), N. Burman (1768), Bonelli (1772), Lamarck (1786), Aiton (1789), Lourero (1790), Redoute (1799), and Willdenow (1799).

**19th, 20^th^, and 21^st^ Century Literature Examined (by region and year):—**

**South Asia and Indian Ocean:—**Roxburgh (1814) [Bengal], Moon (1824) [Sri Lanka], Buchanan-Hamilton (1825) [Northeastern Bengal], Ainslie (1826) [Tamil Nadu], Roxburgh (1832) [India general], Wight (1852) [Tamil Nadu], Bedomme (1872) [South India], Stewart (1874) [Northwest and Central India], Hooker (1875) [“British India”], Trimmen & Hooker (1898), [Sri Lanka], Talbott (1902) [Bombay Presidency], Prain (1903) [Bengal], Cooke (1908) [Bombay Presidency], Burkill (1910) [Nepal], Rama Rao (1914) [Tranvancore], Duthie (1915) [Upper Gangetic Plain and Sub-Himalayan Tracts], Gamble (1915) [Presidency of Madras], Bamber (1916) [Punjab and Pakistan], Parker (1918) [Punjab, Hazara, Delhi], Blatter & Hallberg (1919) [Rajasthan], Haines (1921) [Bihar and Orissa], Parker (1927) [Chakrata, DehraDun, Saharanpur], Kanjilal *et al*. (1940) [Assam], Janse (1953) [India], Saldanha & Nicolson (1976) [Karnataka], Radcliffe-Smith (1982b) [Mascarene Islands], Matthew (1983) [Tamil Nadu], Radcliffe-Smith (1986) [Pakistan], Rao & Sreeramulu (1986) [Andhra], Saxena & Brahmam (1995) [Orissa], Saldanha (1996) [Karnataka], Philcox (1997) [Sri Lanka], Singh (2002) [India], Bhat (2003) [Udupi District, Karnataka], Manjunatha *et al*. (2004) [Davanagere Dist., Karnataka], Eflora of India: *E. neriifolia*, (2007 onwards), Reddy (2009) [Andhra], Binojkumar & Balakrishnan (2010) [India], Ekanayaka *et al*. (2015) [Sri Lanka], Sankara Rao *et al*. (2016), Digital Flora of Peninsular India (2019), Digital Flora of Gujarat (N.D.).

**Southeast Asia:—**Blanco (1837) [Philippines], Miquel (1860) [Sumatra], Kurz (1887), [Myanmar], Warburg (1894) [New Guinea], Hoeck (1896) [New Guinea], Gagnepain (1910) [“Indo-Chine”], Merrill (1918) [Philippines], Merrill (1923) [Philippines], Ridley (1924) [Malay Peninsula], Merrill (1935) [Vietnam], Backer & Bakhuizen (1963) [Java], Radcliffe-Smith (1972) [Thailand], Perry (1980) [East and Southeast Asia], Radcliffe-Smith (1980) [New Guinea], Radcliffe-Smith (1981) [Sumatra], Radcliffe-Smith (1982a) [Central Malaysia], Ho (1992) [Vietnam], Chayamarit & Van Welzen (2005) [Thailand], Esser (2017) [Thailand], DeFilipps (2018) [Myanmar], Pelser (2018) [Philippines], Malaysia Biodiversity Information System (2020) [Malaysia].

**East Asia:—**Bretschneider (1880), He (1997) [China], Ma & Gilbert (2008) [China].

**Non-Asian Countries:—**Haworth (1812), Thonning (1827) [Ghana], Boissier (1862), Karsten (1882), Poisson (1902) [West Africa], Berger (1907), Stapf. (1909) [West Africa], Brown (1913) [Tropical Africa], Croizat (1934), Chevalier (1948), Standley & Steyermark (1949) [Guatemala], Hutchinson & Dalziel (1952) [Tropical West Africa], Wijnands (1983), Burger & Huft (1995) [Costa Rico], Carter (2002) [Africa], Schmelzer (2008) [Tropical Africa].

### Taxonomic treatment of the *Euphorbia neriifolia* complex

#### Euphorbia drupifera

Thonn. (1827: 250); Boissier (1862: 80); Berger (1907: 36). Homotypic synonym: *Elaeophorbia drupifera* (Thonn.) Stapf. (1909: plate 2823); Hutchinson & Dalziel (1952: 65 – 66); Carter (2002: 101); Schmelzer (2008: 238). Type:—GHANA. Without location, *s.d., Thonning 266* (holotype C?, specimen not extant). Neotype:—GHANA. Greater Accra Region: 30 km from Tema Motorway Roundabout on Accra to Aflao Road, Accra Plains, 30m, 25 March 1996, *A. Welsing, M. Merello, & H. Schmidt 36* (BR974659! designated here).

*Euphorbia renouardii* Pax (1902: 61). Type:—Benin [“Dahomey’]. “les environs de Porto-Novo”, *s.d., E. Poisson s.n.* (no holotype or herbarium recorded).

*Euphorbia toxicaria* Afzel. ex Steud. (1840: 615) *nom. nud*.

*Euphorbia elastica* Poiss. & Pax (1902: 60) *nom. provis.* Type:—GUINEA. “near Conakry” s.d., *Poisson s.n.* (no holotype recorded).

**Description:—**Small to medium lactiferous tree up to 22 m with a spreading habit and large rounded crown. Bark grey, branchlets long (50–100 cm), 4–6 sided, becoming terete, with persistent leaf scars, *spine shields* arranged vertically along ridges, obscurely triangular, 5 × 8 mm nearly engulfing leaf scar and flowering eye, greyish tan, with stout stipular spines 2–4 mm. *Leaves* fleshy, obovate, apex rounded or emarginate, 6–28 × 3–10 cm, tapering to a cuneate base and a rounded petiole, 1–2.5 cm, a strong central vein and fine pinnate secondary veins, entire. *Inflorescence:* axillary cymes, usually 3 together, peduncles unbranched or branching once, primary peduncle to 4.5 cm, flaring distally into the involucre, secondary peduncles up to 2.5 cm. *Bracts* broadly deltoid, 7 mm, persistent. *Cyathium* subsessile, 8–12 × 4–5 mm, involucre widely funnel-shaped, *glands* 5, thick, oblong, touching, convex superiorly, dark green becoming brownish yellow, 2.5 × 6 mm, *lobes* 5, greenish, broadly triangular, 2.5 mm with minutely denticulate edge. *Male flowers* numerous, well excerted, approx. 4 mm, jointed, bracteoles variably connate, cuneate, fan-shaped, lacerate, anthers binary, broadly oblong, longitudinally dehiscent, yellow. *Female flower*, stoutly conical superiorly merging into short styles, 1.5 mm, connate to middle, styles bifid. Fruit a drupe of variable size and shape, subsessile, obovate to sub-globose, obscurely 3-lobed, to 5 × 3.5 cm, greenish becoming yellow, endocarp sub-globose to 14 × 11 mm, 1 or 2 ovules usually aborted, single seed oblong with acute apex, 7 × 4.5 mm smooth, greyish brown [Fig. 4].

**FIGURE 4.**
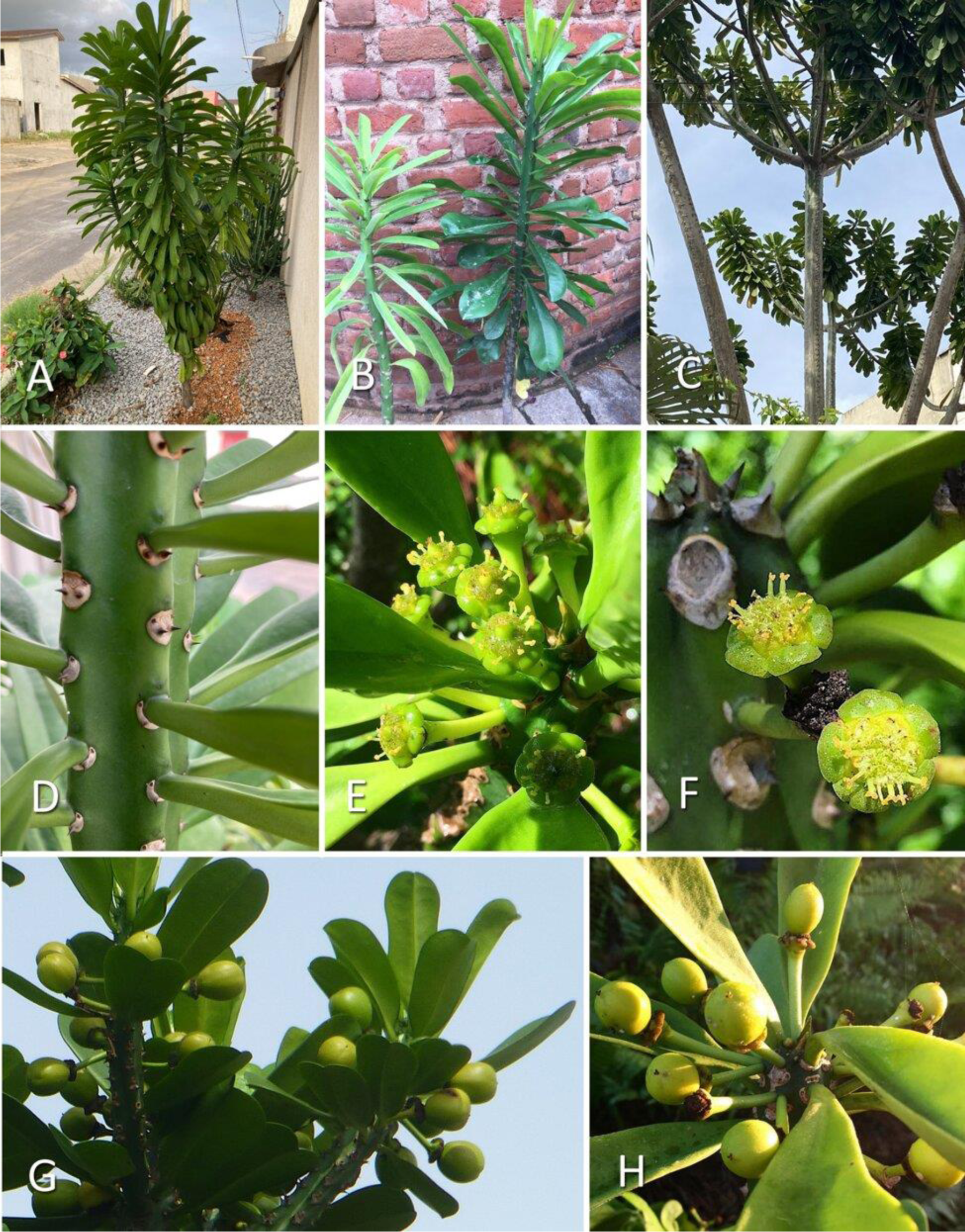
*Euphorbia drupifera* Thonn. (A) Habit of young tree in Abidjan, Côte d’Ivoire, (B) Comparison of *E. nivulia* (left) and *E. drupifera* (right), Tamil Nadu, India, (C) Main branches, Abidjan, (D) Branchlet close-up, Abidjan, (E) Flowering branch, California, USA, (F) Cyathia, top view, California, (G) Fruiting branch Abidjan, (H) Drupes close-up, Florida. Photos A–D, G by Bruce De Jong, E, F, H by Geoff Stein.

**Distribution:—**Tropical East Africa from Guinea west to Uganda and south to Congo (Kinshasa).

**Additional specimens examined:—**CONGO (Kinshasa). Province Orientale: Ituri, near Nduya, by the village Maitatu, marshy area by a river, 11 April 1976, *S. Lisowsky 42398* (BR0000015779946!). COTE D’IVOIRE. Near Bratouedi, 75 km NW of Abidjan, on granite rocks, 100 m, 30 December 1958, *A.J.M. Leeuwenberg 2314* (WAG0084528!). EQUATORIAL GUINEA. Biopo Island, Road from Malabo to Cupappa, 12-13 km, Cocoa Plantation 3 September 1988, *P*. *Carvalho 3607* (P00736498!). GHANA. Adidome, on plains, 10 May 1956, *J.K. Morton A2082* (WAG0084527!). GUINEA. Fouta Djallon, highland plateau between Bitinn and Diaguissa, April 1905, *A. Chevalier 12905* (P00576955!).

**Habitat:—**Forest edges, in rocky areas, flooded coastal plains, occas. in rainforest up to 1000 m.

**Conservation status:—**Probably not vulnerable because of the broad range and ease of growth and cultivation.

**Phenology:—**Flowers and fruit November to May

**Etymology:—**Named for the drupe-shaped fruit.

**Uses:—** Live fence, planted near homes for religious rituals, wood for smoking fish, latex and leaves for treating wounds, ulcers, bites and stings, purgative, and as a fish poison.

**Taxonomic notes:—**Linnaeus first described this African plant under *E. neriifolia* with his rubric “Euphorbium afrium spinosum, foliis latioribus non spinosis” citing Seba (1734: 18, plate 9) who in turn stated he got the specimen from van Beaumont’s garden. We recognise the separation out of this taxon by Thonning, though when he described *E. drupifera* as a new taxon, he did not mention it had been cited by Linnaeus. In 1909, Stapf. proposed the new genus *Elaeophorbia* based on the drupe shaped fruit, and with it the name of this species was changed to *Elaeophorbia drupifera* (Thonn.) Stapf. This has since been returned to the original after molecular genetic studies, thathave shown *Elaeophorbia* to be nested within *Euphorbia* sect. *Euphorbia* (Dorsey *et al*. 2013).

Carter (2002) stated that the type, *Thonning 266*, was not located. The collections manager of Herbarium C of the University of Copenhagen in personal communication confirmed that this type specimen of *E. drupifera* Thonn. has been lost. We selected a specimen from Ghana as a neotype since Thonning’s original collection was from Ghana. This particular specimen clearly shows the character of the leaves, cymes (3 together, and both single and branching forms), cyathia shape, and the sub-sessile drupes.

***Euphorbia ligularia*** (Roxburgh 1814: 36, *nom. nud*.) Buchanan-Hamilton (1924: 285) [*pro parte* excluding *E. neriifolia* while retaining the type of the species]; *Euphorbia ‘ligularia* Roxburgh (1832: 465) *nom. illeg.*, Duthie (1915: 77), Saxena (1995: 1635); Sankara Rao *et al*. (2016: Digital Fl. Eastern Ghats, *E. ligularia*). Type citation: “Habitat in sylvis et ad templae Bengalae orientalis.” Type: INDIA. “Bengal” (without location) *s.d. Roxburgh*, s.n. (see taxonomic notes) (lectotype designated by Chakrabarthi (2019: 3631) [unpublished icons] Icones Roxburghianae [No. 1066 &] 1972 (CAL), remaining original material Icones Roxburghianae [No. 1066 &] 1972 (K!) [Figs. 3 B, 3 C]. Reinstated as a species here, *stat. nov.* (see taxonomic notes below). Heterotypic synonym: *E. undulatifolia* Janse (1953: 69) *syn. nov*. Type:—EUROPE. [without location] *s.d., Janse* (“the collection of the University of Amsterdam”, *A 12501* (lost)).

Misapplied name:

*E. neriifolia* auct. *non* Linnaeus (1753: 451): Cooke (1908: 563); Das (1940: 140); Hooker (1887: 255) *pro parte* (excluding plants now identified here as *E. yadavii* and *E paschimia*); Kurz (1877: 416); Prain (1903: 923); Sankara Rao *et al*. (2019: *Euphorbia neriifolia*) *pro parte* (excluding plants now identified here as *E. yadavii* and *E paschimia*).

**Description:—**A spiny, lactiferous shrub to small tree up to 4 (–6) m with an open-branched, vertically oriented, broadly columnar crown. *Trunk*, lowest branches from base to 1 m above the ground, diameter 10–20 (30) cm, bark light grey and brown, scabrous with persistent scars of teeth. *Branches*, segment length main branches 30–100 cm, width 2.5–3 cm, teeth tall and narrow, sub-confluent to confluent, in five vertical to somewhat spiral rows, tooth height 5 – 8 mm, interval between teeth 2–3 cm, spine shield dark brown with light grey 3 mm leaf scar, shield circular to obovate, with obtuse inferior margin, 3 × 4 mm, spines in pairs, divaricate, 3–7 mm. *Leaves* present on growing segments, partially deciduous during cold season, succulent, somewhat flat and pendulous, entire, borders often undulating when young or very old, obovate, apex rounded to obtuse, ± apiculate, base tapering, cuneate, 10–30 × 3–6 cm, petiole 0.5 cm, central vein pronounced, lateral veins faintly visible, branching at approx. 60–70 degree angle. *Flowers*, single cyme per flowering eye 0–2 mm above spine shield, commonly branched up to three times, *bract* light green, broadly orbicular with irregularly lobed fringe, 4 × 5 mm, extending up to base of glands, faintly keeled, apiculate, *primary peduncle* 2–3 mm, secondary and tertiary peduncles 5–8 mm, central *cyathium* staminate, sessile, very shallow cupular, diameter including glands 8–10 mm, height 4 mm (base to top of lobes), *glands* five, yellowish green, broadly oblong, flattened medially, 4–5 × 2 mm, *lobes* five, 3 × 2.5 mm, orbicular greenish-yellow with irregular lacerate edge. *Male florets* in 5 fascicles of 5(8), often atrophic, indefinite number of spathulate bracteoles of different widths, approx. 2.5–3.0 mm tall, apices highly lacerate and tufted. *Female flower*, pedicel 1.75 mm, ovary sub-ovate, 1.75 × 2 mm, styles connate to distal third then retroflexed, stigma broadened, papillose. Capsules and seeds not seen [Figs. 5, 6].

**FIGURE 5.**
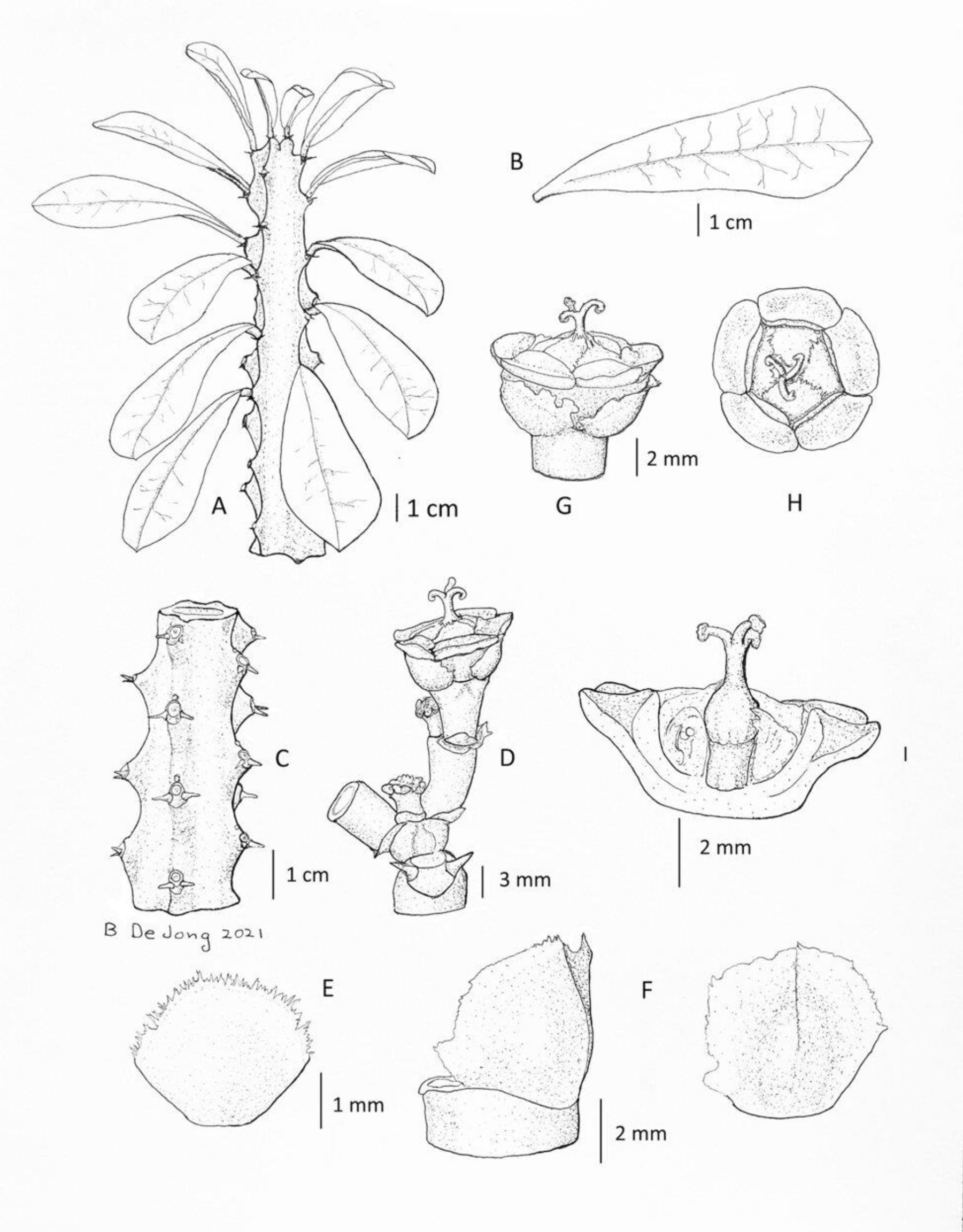
Illustration of *Euphorbia ligularia* Roxb.ex Buch.-Ham. (A) Habit, (B) Leaf, (C) Teeth and spine shields, (D) Cyme with 2 levels of branching, (E) Lobe of cyathium, (F) Bract lateral and front view, (G) Bisexual cyathium lateral view, (H) Bisexual cyathium top view, (I) Bisected cyathium with female flower in situ. Illustration by Bruce De Jong from B. DeJong LGK2a.

**FIGURE 6.**
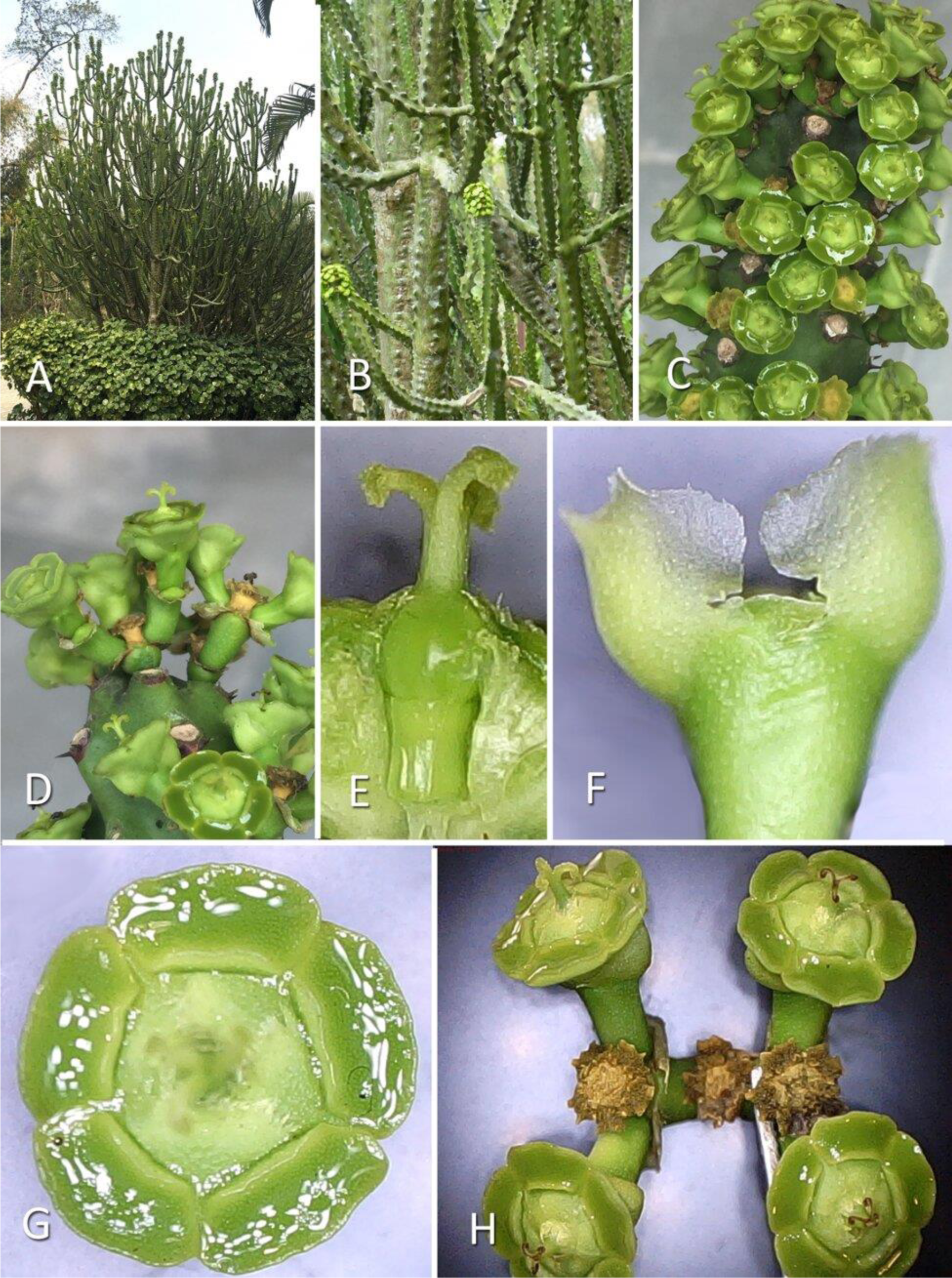
*Euphorbia ligularia* Roxb. ex Buch.-Ham. (A) Habit of topotype plant, (B) Branchlets, (C) Flowering branch, (D) Cymes close-up, (E) Female flower in situ, lateral view, (F) Bracts lateral view, (G) Cyathium top view, (H) Branching cyme. Photos A–H Bruce De Jong.

**Distribution:—**Total extent unknown, but identified in Assam in the districts of Kokrajhar, Chirang, Baksa, Udalguri, Sonitpur, Karbi Angalong, and Gologhat. Roxburgh described it from “Bengal”, which in his day was a vast territory including current West Bengal, Bangladesh, and parts of Assam. Very similar plants are reported in the wild and cultivated in Odisha (Saxena 1995, Sankara Rao *et al*. 2016). [See Map 1.]

**MAP 1.**
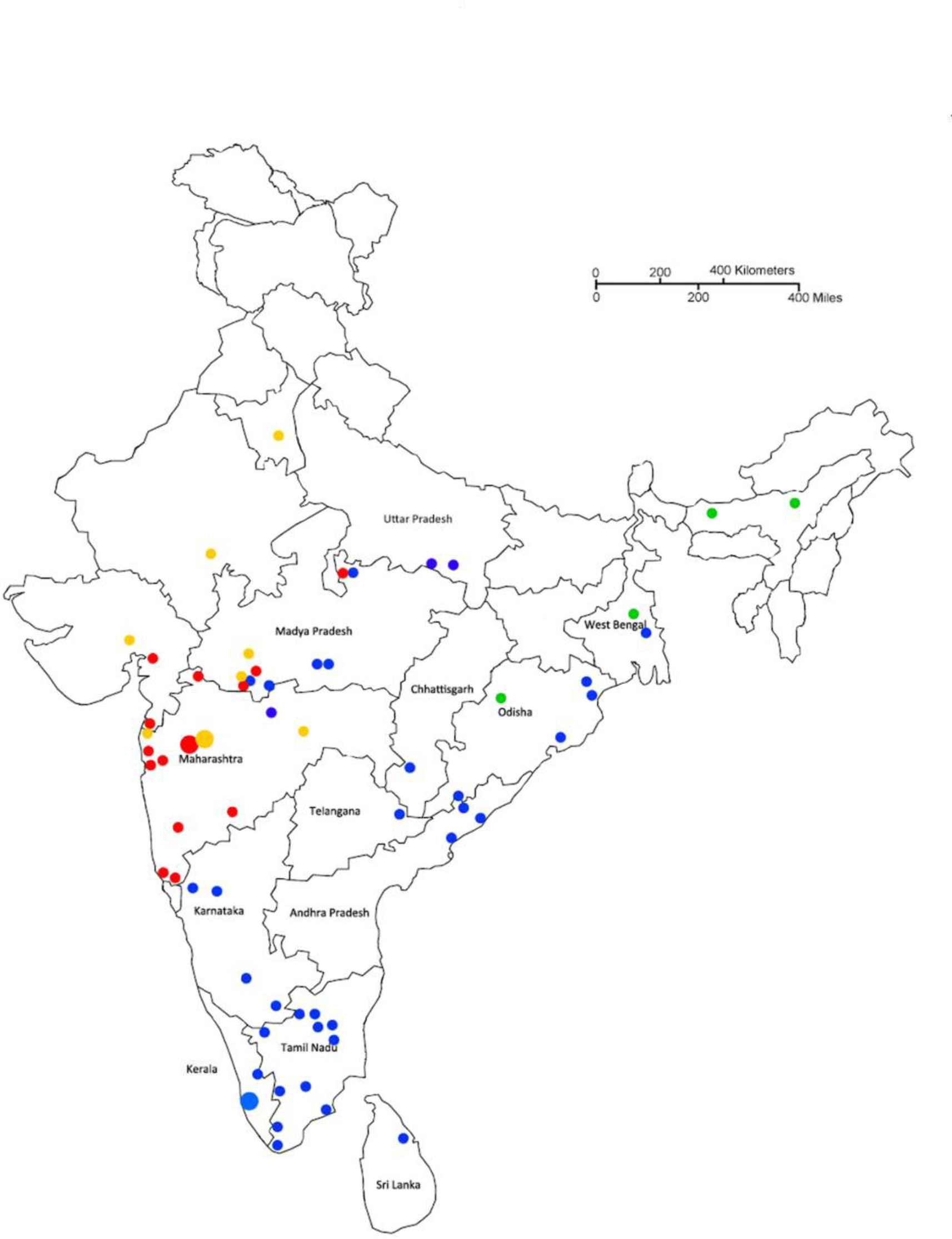
Distribution of South Asian species in the *Euphorbia neriifolia* complex: blue dots *Euphorbia nivulia,* yellow dots *Euphorbia yadavii*, red dots *Euphorbia paschimia*, and green dots *Euphorbia ligularia*. Large dots indicate type locations where known.

**Specimens examined:—**INDIA. Assam: Gologhat District: Kohora, 2 km east of Kaziranga National Park Central Gate at Iora Resort, 16 March 2020, *B. DeJong LGK2* (CAL!)

**Habitat:—**Wild origin unknown, possibly Odisha in Ganjam, Balangir, Bargarh Districts (Sankara Rao *et al*. 2019), but cultivated as sacred plants in home shrines in Assam, especially in the Bodoland Territorial Region, north of the Brahmaputra River and south of the foothills of Bhutan.

**Conservation status:—**Data Deficient, but subjectively only known from cultivation in homes in a limited range in Assam. Fewer people seem to be growing them as the number of home shrines diminishes.

**Phenology:—**Flowers in cold season from February to end of March.

**Etymology:—**The name comes from Rumphius (1743), in which he described the spoon-like shape of the leaves, hence *ligula* in Latin which means small spoon, or tongue. Roxburgh adopted Rumphius’s name (see below).

**Vernacular:—**Bengali: *munsa-sij*, Assamese and Bodo: *sijou*

**Uses:—**As a center piece in Bodo shrines, the five angles and five-lobed flowers symbolising the five aspects of the divinity Bathou.

**Notes:—***Euphorbia ligularia* plants growing in Assam appear to be sterile cultivars, having been propagated by cuttings for many hundreds of years. The lectotype drawing shows male flowers, but no capsules or seeds. Our field study also shows that the plants never have capsules, seeds, or seedlings. Bodo tribal informants informed us that the plants could have come from the forests of Bhutan, but they had not seen them personally. The Bodo people are thought to have migrated across Bhutan from the Tibetan plateau. The plants we found in Assam matched Roxburgh’s drawings closely, both in their use as a religious tree as well as their habit and flower characteristics.

It is also possible that related plants are cultivated in villages and grow wild in Odisha, as per Saxena (1995) who describes a small erect tree 1.8 – 4.5m with tubercles in 5 confluent vertical or slightly spiral ridges, and yellow cymes on very short (3.5mm) peduncles. Photos of a similar plant from Odisha showing the typical sub-confluent flattened tubercles in a saw-tooth pattern and small yellow cyathia on short peduncles are labeled as *E. ligularia* Roxb. on Digital Flora of Eastern Ghats, Sankara Rao *et al*. (2016). More investigation is needed to confirm this.

**Taxonomic notes:—**Even though Roxburgh borrowed the name “*ligularia*” from Rumphius (1743), *E. ligularia* is an Indian species distinct from the Indonesian *Ligularia* of Rumphius which was later named *E. neriifolia* by Linnaeus (1753, 1754).

Roxburgh (1814) first recorded *E. ligularia* R. in the Botanic Garden at Calcutta as a *nom. nud.*, also giving its Bengali name “Munsa-shij” and the date of collection “before 1793”. Under Roxburgh’s supervision, this plant was illustrated and labeled between 1804–1812 (Sealy 1956: 300), Icones Roxburghianae No. 1066 and 1972. Roxburgh brought the manuscript of his *Flora Indica* to England in 1813 with the intent of finishing it there, but died in 1815 (Forman 1997). *Flora Indica* (Roxburgh 1832) was published post-humously, after Buchanan-Hamilton (1824) had published the species.

Buchanan-Hamilton, who succeeded Roxburgh as the Superintendent of the Botanic Garden at Calcutta in 1814 (JSTOR 2013) described *E. ligularia* in 1824 and attributed the name to Roxburgh. Since this is the first validly published instance of *E. ligularia*, according to article 6.3 of the ICN (Turland *et al*. 2018) it takes precedence over the more complete description of the same taxon based on the same type by Roxburgh (1832). Even though *E. ligularia* Roxb. is superfluous and is a *nom. illeg*., the description of the species in Roxburgh (1832) is detailed and complements the lectotype.

However, Buchanan-Hamilton (1824) clubbed together the new *E. ligularia* from Bengal with *E. neriifolia* L. from Indonesia which was growing in European gardens at the time. Since they are morphologically distinct, we designated only the Bengal plant (*pro parte)* as type, following Buchanan-Hamilton’s own type citation.

Boissier (1862:79) was the first to make *E. ligularia* a *syn. nov.* of *E. neriifolia* L. The following authors also supported this view: Beddome (1872: 216), Hooker (1887: 255), Stewart (1874: 439), Parker (1918: 446), Binojkumar & Balakrishnan (2010: 310), and Esser (2017: 28). Govaerts *et al*. (2002) considered it a species. Saxena (1995: 1635) and Rao *et al*. (2016) also treated it as a species.

We have reinstated *E. ligularia* because of its many unique characteristics and specific geography which makes it distinct from *E. neriifolia*.

Binojkumar & Balakrishnan (2010: 312) considered Roxburgh’s Icones No. 1972 in CAL to be lectotype, but this was not validly designated. Chakravarthi (2019: 3631) validly designated this same original material as lectotype.

*E. undulatifolia* Janse (1953: 69) was described based on plants in the author’s personal collection and those found in the Botanic Gardens of Brussels. The author presented photos and descriptions, showing these plants to be morphologically distinct from *E. neriifolia* and *E. nivulia.* The main distinguishing feature is the undulating borders of the leaves. Correspondence with the Naturalis Biodiversity Center which now houses the former University of Amsterdam collection failed to locate any record of the type specimen designated in the protologue in Leiden or in Amsterdam.

The Director of the *Euphorbia* section of the National Botanic Garden of Brussels in Meise provided us with photos of plants in their collection bearing the name of *E. undulatifolia* which match the images in the protologue. The sub-confluent teeth and drooping leaves appear identical to those of *E. ligularia*. Undulating borders of leaves, though not constant, are also seen with new growth and old leaves of the *E. ligularia* plants from Assam. The plants have not bloomed in Meise. A literature and internet search failed to show any other collections of *E. undulatifolia*.

***Euphorbia neriifolia*** Linnaeus (1753: 451) *pro parte* [excluding *E. nivulia* Buch.-Ham. and *E. drupifera* Thonn., while retaining the type of the species]; Osbeck (1757: 205, 209), N. Burman (1768: 111); Bonelli (1772: t. 28); Houtyn (1777: 738); Lamarck (1788: 415); Lourero (1790: 298); Redoute (1799: plate 46); Willdenow (1799: 884); Haworth (1812: 130); Boissier (1862: 79); Ridley (1924: 182); Berger (1907: 34); Gagnepain (1910: 239); Burkill (1935: 979); Merrill (1935: 242); Standley & Steyermark (1949: 107); Janse (1953: 70); Wijnands (1983:100); Radcliffe-Smith (1980: 185); Radcliffe-Smith (1982a: 19); Radcliffe-Smith (1982b: 93); Corner (1988: 291); Burger & Huff (1995: 119); Ma (1997: 60, plate 15, fig. 1-3); Esser (2005: 290); Esser (2017: Euphorbia); Ma & Gilbert (2008: 300, plate 352, figs. 1-3); Malaysia Biodiversity Information System (MyBIS) (2020: *E. neriifolia*); Pelser (2020: *Euphorbia* L.). Homotypic synonym: *Elaeophorbia neriifolia* (L.) Chevalier (1948: 348).

Type citation: “Habitat in India.” Type:—(lectotype LINN 630.1! designated by Radcliffe-Smith (1982b), Epitype (designated here):—INDONESIA. East Java Province: Pasaruan Regency, Bangil Division, 6 July 1925, *C.A. Backer 37520* (L. 2225125!) [Figs. 2 D, 3 D]

*Euphorbia edulis* Loureiro (1790: 298); Boissier (1862:80); Gagnepain (1910: 239 – 240, 254) [as prob. var.]; Merrill (1935: 242) [as *syn. nov*.]; *Tithymalus edulis* (Lour.) Karsten (1882: 587). Type citation: “Habitat culta in hortis Cochinchinae”, (no collection data given).

*Euphorbia pentagona* Noronha (1790: 14) *nom. nud*.

*Euphorbia pentagona* Blanco (1837: 413) *nom. illeg.* non Haworth (1828: 187)

*Euphorbia complanata* Warburg (1894: 196), *syn. nov.*, Hoeck (1896: 234). Type:— NEW GUINEA. Morobe Dist.: Finschhafen, on hills behind station, 5 May 1889, *Hellwig* 678 (not extant). Neotype:— NEW GUINEA. Morobe Dist.: Markham Valley, Kajabit [Kaiapit], Mission grounds, 300 m, 25 July 1939, *Clemens* 10495, (neotype MICH1283947! designated here, isoneotype: L2225652!)

Misapplied names:

*Euphorbia ligularia* auct. mult. *non* Buchanan-Hamilton (1824: 245): Miquel (1859: 418); Backer & Bakhuizen (1964: 501); Radcliffe-Smith (1972: 265); Radcliffe-Smith 1981: 296); Ho (1992: 358, fig. 4663).

**Description:—**Much branched, armed, lactiferous, succulent shrub to small tree 4–6 (–7) m high, with upwardly curving branches that form a densely packed globose crown. *Trunk*, lowest branches 50 to 150 cm above the ground, 10–20 (–30) cm wide, bark light grey with smooth scales and persistent scars of teeth, somewhat fissured dark grey in very old trees. *Branches* whorled, segment length main branches 10–50 cm long, terminal branchlets 10–20 cm high, 2.5–3.5 cm wide, teeth, narrow, sub-confluent in five spiral rows, teeth 1–4 mm long, interval between teeth 2–3 cm long, spine shield dark brown, obovate, with obtuse inferior margin, 3–4 × 5–6 mm, spines in pairs, divaricate, 2–5 mm long. *Leaves* bunched near the growing ends of branches, deciduous, spoon-like concave up, spathulate, glabrous, succulent, entire, oblanceolate or elongated obovate, 6– 15 × 2.5–4 cm, apex obtuse, apiculate, petiole 0.5–1cm long, central vein pronounced, lateral veins faintly visible, branching at 70–90 degree angles. *Flowers* in single cyme per flowering eye, 0–2 mm above spine shield, *bract* pale green, broadly obovate, 7 × 7 mm, incorporating lateral edge of glands, entire, keeled, apex obtuse and apiculate, primary *peduncle* 3–8 mm long, secondary peduncle 8–12 mm long with buds present for a tertiary peduncle, central cyathium staminate, sessile, very shallow cupular, 8–10 mm in diameter, 4.5 mm (base to top of lobes) high, glands five, reddish brown to red turning greenish with maturity, convex, oblong, 4-5 × 1.5 mm with wavy margin, lobed, lobes 5.3 × 2.5 mm, erect, orbicular greenish-yellow with irregular lacerate yellowish fringe. *Male florets* in 5 fascicles of 5–7 flowers, progressively maturing from centre to periphery, pedicel off-white, hyaline, ca. 2.5 mm long, indefinite number of spathulate bracteoles of different widths, 2.5–3.0 mm tall, apex highly lacerate and tufted, stamen filament pale red ca. 1.5 mm long, anthers flattened and oblong, ca. 0.7 mm long, red, longitudinally dehiscent. *Female flower*, pedicel 2 mm long, ovary ovoid, ca. 3 × 2 mm, 3-lobed, styles connate to below center, stigma broadened, papillose. Capsules and seeds not seen. [Figs. 7, 8].

**FIGURE 7.**
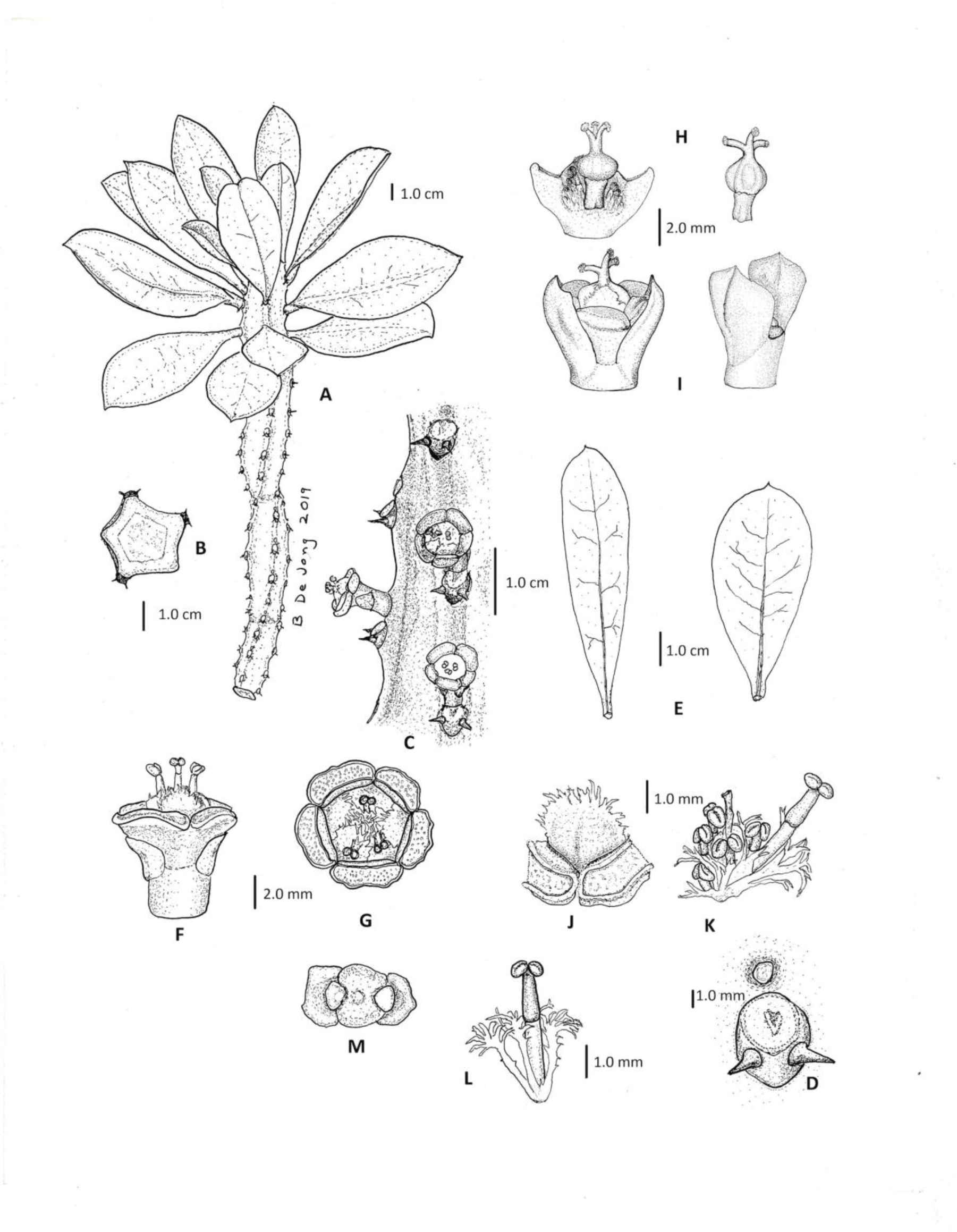
Illustration of *Euphorbia neriifolia* L. (A) Habit, (B) Cross-section, (C) Teeth and spine shields close-up, (D) Spine shield close-up, (E) Leaves, (F) Staminate cyathium lateral view, (G) Staminate cyathium top view, (H) Female flower lateral view, (I) Bracts in situ, (J) Lobe of cyathium, (K) Fascicle of male flowers, exposed, (L) Male flower with bracteoles, (M) Bracts surrounding buds of peduncles, top view. Illustrated by Bruce De Jong from Amed Beach, Bali.

**FIGURE 8.**
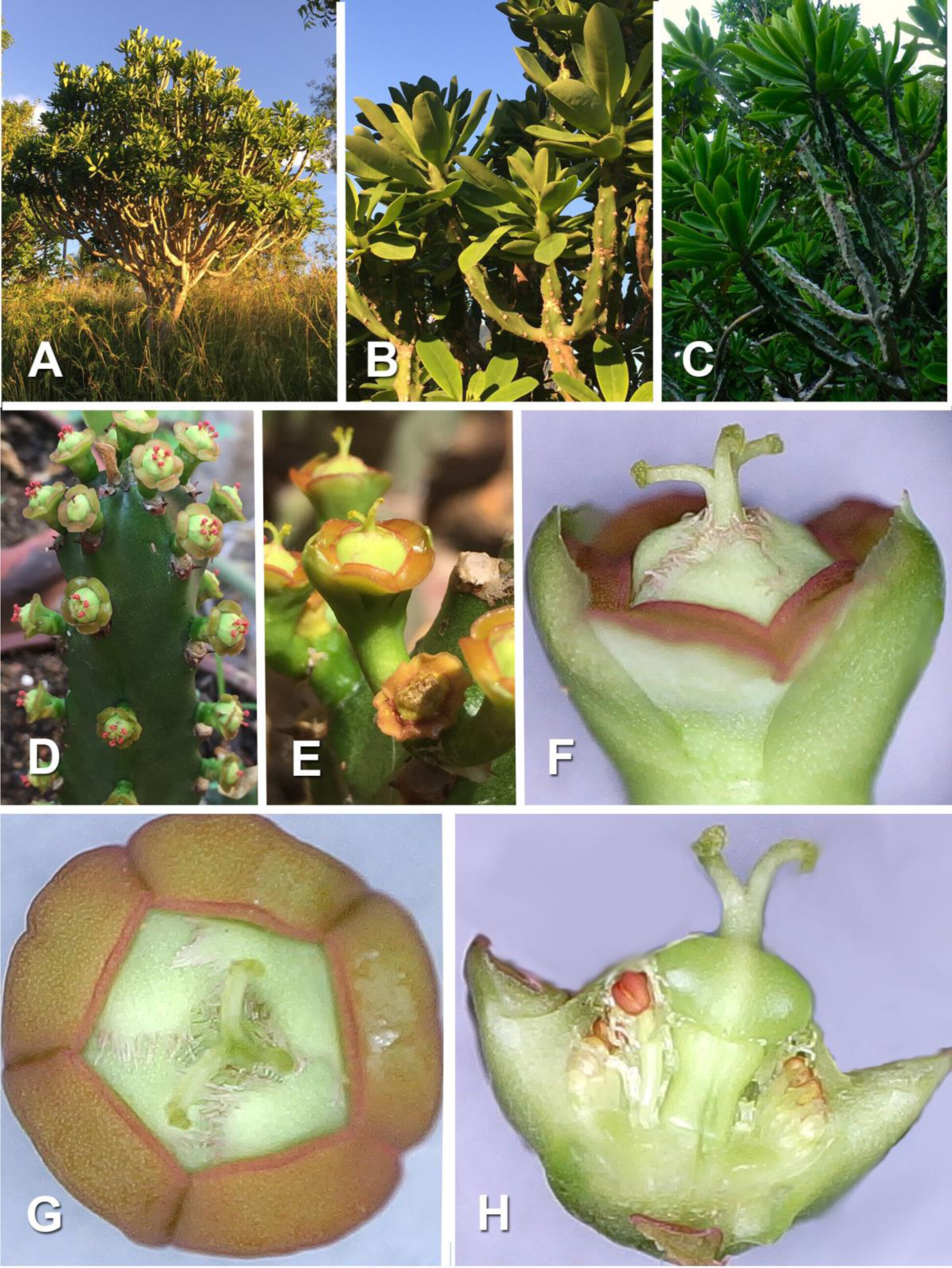
*Euphorbia neriifolia* L. (A) Habit of isotype plant in the field, (B) Leaves, (C) Branchlets, (D) Flowering branch, (E) Simple cymes, (F) Bisexual cyathium lateral view, (G) Bisexual cyathium top view, (H) Bisected cyathium showing male and female flowers. Photos A–H Bruce De Jong.

**Phenology:—** According to Rumphius, only blooms after a long drought. Bali specimens bloomed in the cold, dry season (February–March), Flora of China (June– September). Data from herbarium sheets with flowers: Vietnam (March), New Guinea and Java, dry season (July–August), Caribbean (November–May).

**Distribution:—**Wild origin obscure, possibly native at least in NE New Guinea (Warburg 1894). An ornamental, mainly planted in China, Indonesia, Malaysia, New Guinea, Philippines, Thailand, Vietnam, and several Caribbean and Central American countries.

**Habitat**:—Wild origin obscure. Commonly planted in hedges and cultivated in gardens around the world.

**Conservation Status:—**Considering its popularity as a garden plant and ease of reproduction by stem cuttings, IUCN Redbook 2022 conservation status would be LC or “least concerned”, although if fertile varieties were found in the wild, this would have to be reevaluated.

**Additional specimens examined:—**CHINA. Guangdong Province: Zhangjian (Kouy-Tcheou), Nov. 1906, *M. Cavalarie 3098* (P00702747!). Hainan: by side of river, 2 Jan. 1934, *C. Wang 36132* (NY04048528!); in open waste places, 12 Jan. 1934, *H.Y. Liang 64536* (NY04048529!). CUBA. Villa Clara Province: Santa Clara, escaped from fence rows to pasture, vicinity of Soledad, June 1941, *R.A. Howard 5407* (NY01428980!). GUATEMALA. 7 Oct. 1949, *P.C. Standley 74183* (F1090882). GUYANA. Region 4 – Demerara-Mahaica: Georgetown, Garden, July 1923, *A.C. Persaud 382* (F532936!). Region 1 –Barima Waini: Barima River, Koriabo house yards, 2 Nov. 1996, *T. van Andel, N. Roberts, G. Ford & N. George 1281* (U0065341!). HAITI. Department du Nord: north of St. Michel de l’Atalaye plantation, 20 Nov. 1925, *E.C. Leonard 7300* (NY01428979 !). HONDURAS. 30 Oct. 1948, *P.C. Standley 13886* (F1387547). INDONESIA. West Java: Bogor, Bogor Botanical Garden, 1922, VIII. H. *16*, (L2225632!). North Malukku: Halmahera, Galela, 25 m, 22 Oct. 1921 ?. *Beguin 1832* (L0272683!). Papua Province: Merauke, *s.d. J.W.R. Koch 1904* (L0151586!). East Nusa Tenggara, Manggarai, 20 April 1978, 600 m, *Fr. E. Schmutz 4210* (L0272681!). West Sumatra: Solok Regency, Tigo Lurah, Simanau, 28 Nov. 2012, *HRN & RPN 51*, TAMA130020020306, (also known as ANDA00012544!). JAPAN. Okinawa Prefecture: Ryukyu Islands, Miyako-Jima, 30 Aug. 1956, *R.L. Hetzler 1* (L0147375!). MARTINIQUE. Cultivated in Martinique and Guadeloupe, June 1901, *Fr. A. Duss 4054* (NY01428981!). MONTSERRAT. St. Anthony Province: Plymouth, in a yard, *R.A. Howard 19250* (NY01428977!). NICARAGUA. Departamento de Chontales: Rio Las Vainillas, road to Santo Tomas-Juigalpa, cultivated, 200 m, 21 Jan 1983, *J. C. Sandino & S. I. Martinez 3890* (MO3190022!). PAPUA NEW GUINEA. Morobe: Kalanza, hot grassy hills, always on coral rock, 1800 – 2000 feet, 28 Dec. 1937, *M.S. Clemens 7931* (B101123376!, B101123377!); Markham Valley, Kaiapit (Kajabit), Markham valley, Mission grounds, 300 m (800-1000 feet), 25 July 1939, *M.S. Clemens 10495*, (MICH1283947!, L2225652!). PHILIPPINES. Luzon: Manila, Jan. 1915, *E.D. Merrill 823* (L0272682!), THAILAND. Chiang Mai Province: cultivated 300 m, 9 Jan. 1921, *A.F.G. Kerr 469* (L0151582!). TOBAGO. Tobago Botanic Station, 22 Feb. 1910, *W.E. Broadway 3503* (L2225631!), (NY01428976!). TRINIDAD. San Fernando, cultivated in gardens, 15 April 1874, *O. Kuntze 963* (NY01428978!, P05589645!). US VIRGIN ISLANDS. St. Croix: Anna’s Hope, planted, 5 May 1924, *J.B.Thompson 751* (NY01428983!). VIETNAM. Ben Tre Province: Ba Tri District (In the forests of Tri Huyen), March 1872, *L. Pierre 1083* (P00702755!). Ha Nam Province: Thinh Cao, growing in fence lines, 10 April 1883, *H.F. Bon 2055* (P00702753!). Lao Cai Province: Bac Ha District, Bao Nhai, 8 Dec. 1935, *E. Poinlane 24989* (P00702748!).

**Etymology:—**The specific epithet “Nerii folio” was first published in Breyne (1689) and adopted by Linnaeus (1753), indicating that the leaves resemble those of *Nerium* L. or Oleander.

**Vernacular names:—**Banda Islands: *carambau dinar*, China: *fu yong fa*, *que haan, jin gang zuan*, E. Nusa Tenggara: *bilas,* Java: *daun sudu sudu*, Malaysia: *sesudu*, Bali: *blatung bun* “spoon cactus”, Maluku islands: *pfuggi lida*, Myanmar: *sha-soung,* Philippines: *bait, soro-soro*, *sorog-sorog*, Thailand: *som chao,* Vietnam: *xuong rong, giang lam, laong*.

**Uses:—**Planted as hedge, ornamental; leaves boiled with syrup and eaten, used as fish poison, vermifuge, cathartic, emetic, diuretic, asthma, applied for earache, warts.

**Notes:—**It appears that *E. neriifolia* rarely produces fruits. The only fruiting evidence is recorded in the *Flora of China* (Ma & Gilbert 2008 fig. 352) and in the protologue of *E. complanata* from Papua New Guinea. It is possible that centuries of propagation by cuttings has produced sterile cultivars, similar to *E. trigona* and *E. lactea*. No records of juvenile plants were found in the literature. Since the earliest and most pervasive records are from East and Southeast Asia, this region would be the most likely place of origin.

**Taxonomic considerations:—***Euphorbia neriifolia* can only apply to the plants that match the lectotype and the C. Commelin plant cited in Linnaeus’s protologue, namely the form widely planted in Indonesia and China. The lectotype’s presumptive collection details in 1752 are drawn from Osbeck’s travel diary in which he records the date and place he encountered *E. neriifolia*. (Osbeck 1757).

Our study of the literature makes it clear that the only Indian plant cited in the protologue of *Euphorbia neriifolia* that could have been growing in the Europe at the time of Linnaeus was *E. nivulia* (“*Ela-calli”*), obtained from Sri Lanka and/or from Malabar (Kerala), India. There are no illustrations or references to collections of other plants of this complex from India during this period. At the time, plant collectors were often clergy, traders, or colonial officials and had only a superficial access to coastal trading posts such as Surat, Bombay, Goa, Cochin, Madras and Calcutta (Stearn 1957).

Regarding Linnaeus’s “Habitat in India” in the protologue, this could apply nonspecifically to “*Ela calli*” as well as to plants from Indonesia and China. Linnaeus’ sense of geography was known to be imprecise. “It seems that Linnaeus had a very confused idea with respect to the position of China, and we cannot but think, that he considered the latter name to be a synonym for India.” Bretschneider (1857: 90). This reference to India has been a source of great confusion.

Linnaeus’s original description of the taxon was very general: a spiny lactiferous, succulent shrub to small tree with succulent obovate to lanceolate leaves bunched near the ends of branches, with spines arranged in five spiral rows of tubercles along the branches. Because of this lack of specificity and Linneaus’s reference to India, several centuries of authors have assumed that all the South Asian plants which share these general characteristics are conspecific with the plants from Indonesia and China. Our field observations of plants from across India, both wild and in cultivation, have failed to find any plants that match the characteristics of the Indonesian and Chinese plants of *E. neriifolia*. Therefore, the name is misapplied to similar South Asian plants which share the general spiral teeth pattern and bunchy leaf habit.

Keeping this in mind, we went through the Indian literature and herbaria sheets and assigned plants labeled as *E. neriifolia* or *E. nivulia* to their respective taxa wherever possible, in particular *E. nivulia*, *E. ligularia*, *E. paschimia*, and *E. yadavii*. In some cases, different authors had clubbed two or more taxa together, and this is indicated as “*pro parte*”. In other cases, insufficient details were given to make a judgement and we excluded these.

The lectotype of *Euphorbia neriifolia* bears demonstrably ambiguous shriveled leaves and no flowers and therefore cannot be used to apply the name precisely. Hence, following Art. 9.9 of the ICN (Turland *et al*. 2018) we proposed an epitype from Java with well-preserved cyathia for the more precise application of the name. Even though this sheet is labeled *E. ligularia* Roxb., the plant is clearly *E. neriifolia* based on morphology and geography.

Miquel (1859), Backer & Bakhuizen (1964), Radcliffe-Smith (1972, 1981), and Ho (1992) while describing plants in Southeast Asia that match the type of *E. neriifolia* used the name *E. ligularia* Roxb. which should only apply to the plants from the Northeast India. In addition, Cook (1908) used *E. ligularia* to describe plants from the Bombay Presidency which are different from the type of *E. ligularia,* hence our designation of *Euphorbia ligularia* auct. mult. *non* Roxb.

Warburg (1894:196) described a plant growing in the hills “behind the station” near Finschhafen in NE Papua New Guinea which he called *Euphorbia complanata* Warb. Though he acknowledged that it is very similar to *E. neriifolia*, he felt that it was a separate species because the leaves which aggregate near the branch tip are flattened and much longer. He also felt that because it was growing in the wild and had fruits and seeds, it could well be native to the region. Though Warburg designated a holotype (Hellwig no. 678), no such specimen could be found in JE or B where many of Hellwig’s specimens were preserved. The curator of JE informed us that many specimens in B were destroyed during WWII. Since no other original material could be traced, we designated a neotype collected by Clemens from the same region in 1939, a specimen that demonstrates both the habit and cymes clearly. By our analysis, *E. complanata* cannot be distinguished from other flowering *E. neriifolia* specimens and thus is conspecific.

***Euphorbia nivulia*** Buchanan-Hamilton (1825: 286); Boissier (1862:79); Wight (1852: t.1862); Beddome (1872: 216); Stewart (1874: 439); Kurz (1877: 447); Hooker (1887: 255); Talbot (1902: 207); Prain (1903: 923); Brandis (1906: 558); Berger (1907:35); Rama Rao (1914: 352); Bamber (1916: 85); Parker (1918: 446); Haines (1921: 242); Gamble (1925: 1277); Parker (1927: 415); Das (1940: 140); Razi (1946: 53); Janse (1953: 71); Ramaswamy & Razi (1973: 342); Saldanha & Nicolson (1976: 338); Matthew (1983: 1435); Wijnands (1983: 100); Radcliffe-Smith (1986: 118); Rao & Sreeramulu (1986: 423); Saxena (1995: 1635); Saldanha (1996:139); Singh (2002: 51); Manjunatha *et al*. (2004: 354); Balakrishnan & Chakrabarty (2007: 276); Eflora of India (2007 and onwards: *Euphorbia nivulia*); Reddy & al (2009:182); Binojkumar & Balakrishnan (2010: 315); Ekanayake *et al*. (2015: 30); Sankara Rao & al (2019); Digital Flora of Gujarat (N.D.). Type citation: “Habitat ubique in Indiae sepibus.” Type:— (lectotype Fig. 43 of Rheede (1679: 83) “Ela Calli’ designated by Wijnands 1983: 101). Heterotypic synonyms: *E. helicothele* Lemaire (1857: 100). Type—?Madagascar, n.d. n.s. “Elle a été introduîte de Madagascar…et par les soins zêlès du même M. Richard.”, but description is most probably of *E. nivulia*; *E. nivulia var. helicothele* (Lem.)

Boissier (1862: 80). Type**—** “In Madagascaria (Richard!)”. Misapplied names:

*E. neriifolia* auct. *non* Linnaeus (1753: 451); Miller (1768: Euphorbia No.5); Aiton (1789: 134); Roxburgh (1832: 467); Miquel (1859: 418); Cook (1908: 564); Duthie (1915:76); Philcox (1997:192).

**Description:—**Small lactiferous, succulent tree up to 7 (–10) m, irregular hemispheric crown, crowded, straight branches, often in whorls. *Bark* dark brown, deep vertical fissures when mature. *Branches* dull green becoming grey-brown, spreading, long, straight to sl. curving, terete with tubercles up to 3 mm long obscurely arranged in 5 spiral rows. *Spine shields* dark brown, circular, 6 × 6 mm, with thick divaricate spines in a pair up to 5 mm, prominent leaf scars covering most of shield. *Leaves* succulent, crowded at the tip of branches, 10–25 cm × 3–8 cm, bright green above, glaucous below, entire, sl. concave upwards, obovate with rounded, retuse apex, or oblanceolate to oblong with obtuse mucronate apex, petiole 1.5–3 cm, base cuneate, prominent central nerve, approx. 8 pairs of faint lateral nerves, deciduous during the dry season. Single *cyme* from flowering eye, 1-2 mm above spine shield, primary, secondary, and tertiary peduncles 10–25 mm, *bracts* broadly ovate 3 × 3 mm, greenish quickly becoming brown and scarious, central *cyathium* staminate, lateral cyathia bisexual, usually one branching but up to two in mature trees, involucre shallow cupular, 10–12 × 4–5 mm, glands oblong, greenish yellow, 4–5 × 2–2.5 mm, lobes broadly oval, lacerate edges, 4 × 3 mm, greenish-yellow. *Male flowers* in 5 fascicles of 6–8 flowers, pedicel faintly greenish 3.5– 4 mm, filament red 1.5–2.0 mm, anthers dark red in a pair, reniform, 1 mm, fissure longitudinal, bracteoles narrow spathulate, highly lacerate, 0.5–1.5 mm. *Female flower* pedicel approx. 1 mm, ovary subglobose with 3 lobes, 3 × 3 mm, styles connate to middle, bifid at tips, stigmata papillose, *capsule* excerted from the involucre, pedicel of capsule retroflexed when immature becoming curved or straight, approx. 10–15 mm, capsule, dark green to reddish, 10–15 × 5–7 mm, deeply lobed, cocci compressed distally, prominent suture between cocci, *seeds* globose, 3.5 × 3.5 to 4 × 4 mm, brown with tan spots and a tan zone surrounding the dark raphe. [Figs. 9, 10].

**FIGURE 9.**
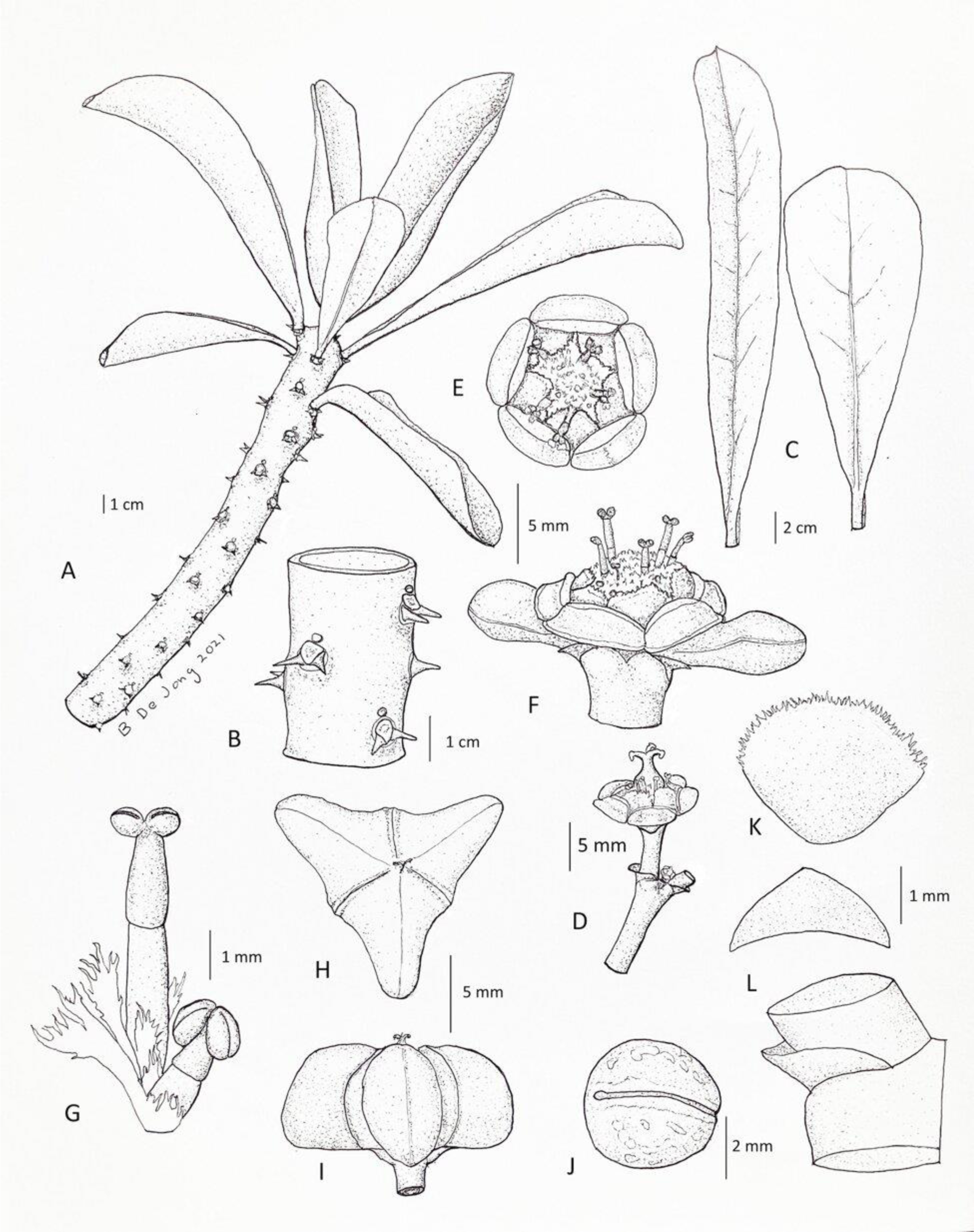
Illustration of *Euphorbia nivulia* Buch.-Ham. (A) Habit, (B) Stem close-up, (C) Leaves, (D) Simple cyme with bisexual cyathium, (E) Staminate cyathium top view, (F) Staminate cyathium lateral view, (G) Male flowers with bracteoles, (H) Capsule top view, (I) Capsule lateral view, (J) Seed raphe view, (K) Lobe of cyathium, (L) Bracts front and lateral views. Illustrated by Bruce De Jong from Kalrayan Hills.

**FIGURE 10.**
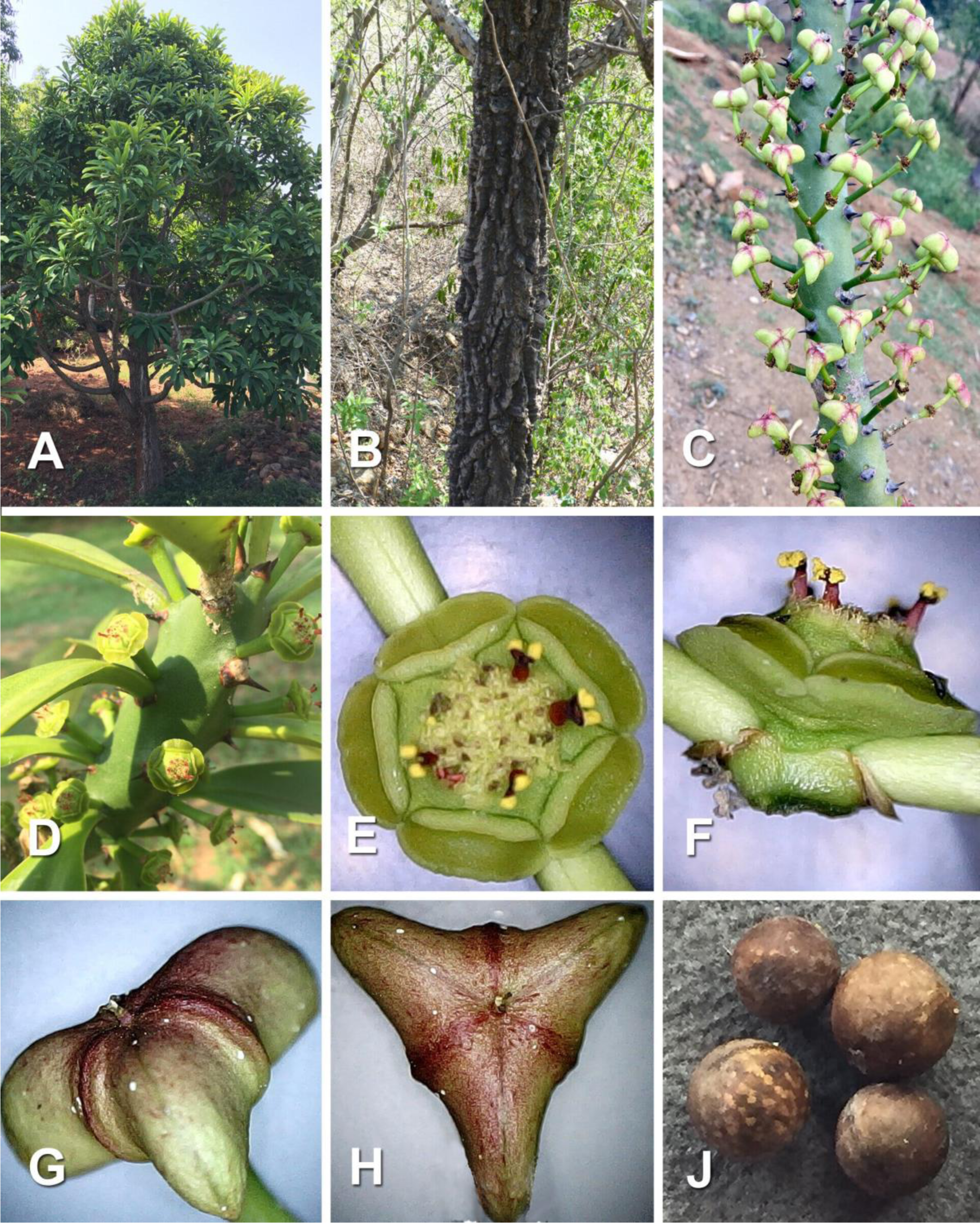
*Euphorbia nivulia* Buch.-Ham. (A) Habit, (B) Trunk, (C) Fruiting branch, (D) Flowering branch, (E) Staminate cyathium top view, (F) Staminate cyathium lateral view, (G) Capsule lateral view, (H) Capsule top view, (J) Seeds. Photos A–J Bruce De Jong.

**Distribution:—**India (Tamil Nadu, Kerala, Karnataka, Andhra Pradesh, Telangana, Maharashtra, Madhya Pradesh, Uttar Pradesh, Chhattisgarh, Odhisa, West Bengal), Sri Lanka, and Myanmar [See Map 1].

**Specimens examined:—**INDIA. Andhra Pradesh: East Godavari Dt., Ontimamidi Vagu Thed Palandi, Jamladugu, 27 March 1988, *N. Rama Rao & T. Ravishankar 86546* (MH!). Kadappa Dt., Palkonda Hills, 1500 ft, July 1884, *J.S. Gamble 14028* (CAL!). Vishakhapatnam Dt., Kilagada, 9 March 1965, *G.V. Subbarao 22577* (MH!); Jalaput, 7 January 1969, *G.V. Subbarao 33306* (MH!); Simhachalam Hill eastern slope, 6 May 1964, *G.V. Subbarao 19379* (MH!). Chhattisgarh: Bijapur Dt., Kandla Patri-Indravati Tiger Reserve forest, 20 May 1987, *A. Kumar 16278* (CAL!). Karnataka: Bagalkot Dt., Badami, 8 March 1909, *S. V. Shevare s.n.* (BSI!). Hassan Dt., Nagpuri Forest, Arasikere, 1 April 2015, *R.K. Swamy & A. Singh 0694* (JCB!). Mandya Dt., Shivasamudram, in dry deciduous forest, 3 March 1978, *S.M. Ahamed* KFP324 (JCB!). Shivamogga Dt. [Mysore], Talguppa, October 1908, *A. Meebold 7305* (CAL!). Kerala: Trissur Dt., Solayar to Malakkampara ghat road, 1300m, 5 February 1984, *K. Ramamurthy 80846* (CAL!). Madhya Pradesh: Damoh Dt., Damoh, 2 October 1908, *J.R. Parno 29935* (CAL!). Hoshangabad Dt., Dhupgarh, 430m, 20 August 1949, *V. Narayanabirami 3380* (CAL!). Nimar Dt., 7 December 1907, *D.O. Witt* 27835 (CAL!). Sagar Dt., Hirapur, 450m, 3 March 1960, *K. Subramaniyam 10171* (CAL!); Sagar Cantonment, 19 Oct. 1906, *A. Blythe 28728* (CAL!). Maharashtra: Akoli Dt., Patur, Medshi village, 21 February 1977, *S.Y. Kamble 152711* (BSI 106033!); Odisha: Khorda Dt., Chandaka Forest, 24 Feb. 2016, A*. Singh, R.K. Swamy, N. Page & K. Sankara Rao 0470* (JCB!). Mayurbhanj Dt., Simlipal Forest, Baniabasa, 23 March 2002, *D.D. Bahali & D.K. Agarwala 560* (CAL!). Tamil Nadu: Dharmapuri Dt., Hogenakkal, 14 March 1965, *E. Vajravelu 23543* (MH!); Dharmapuri to Jhalwania, 10 February 1987, *M. Prasad 39455* (CAL!). Kanyakumari Dt., Poovathu Odai, Boothapandy Range via Karumparai Palrulum, 23 February 1983, *A.N. Henry 77127* (CAL!). Nilgiri Dt., Mayar River bank, 16 February 1972, *B.D. Sharma 39809* (MH!). Ramnad Dt., Aiyanar Kovil forest, 12 March 1970, *E. Vajravelu 33683* (MH!). Salem Dt., Chinnakalrayans, 700 m, 20 February 1979, *T.S. Jeyaseelan 21871* (RHT!). Yercaud, 23 March 2016, *R.K. Swamy, N. Page, A. Singh & K.S. Rao 361* (JCB!). Tenkasi Dt., Honey Falls to Courtallam, 20 March 1958, *K. Subramanayam 5598* (CAL!). Tiruchirapalli Dt., Pachaimalais, 1000 m, 23 March 1977, *D.I. Arockiasamy 7004* (RHT!); Pachaimalais, 100 m, 29 March 1983, *K.M. Matthew, S.J. Britto, & N. Rani 29409* (RHT!). Telangana: Bhadradri Kothagudem Dt. [Godavari], Dummagudem 500ft., February 1885, *J.S. Gamble 15196* (CAL!). Uttar Pradesh: Mirzapur Dt. Kotwa, 15 March 1970, *G. Panigrahi 12666* (BSA!); Prayagraj (Allahabad) Dt., Mahuli Kala, 27 April 1971, *G. Panigrahi 11259* (BSA!) West Bengal: Burdwan Dt., n.d., *A.J. Dutt 1163* (CAL!). SRI LANKA. Anuradhapura Dt: Padaviya, Wahalkada Saddle Dam, rocky dry zone forest, 17 August 2014, *S.P Ekanayaka Y14* (PDA!).

**Habitat:—**Dry deciduous zone, rocky slopes and edges of forests 400–800 (1000) m, also scrub jungle, occasional.

**Conservation status:—**Our subjective impression is that the species is vulnerable, being found in fairly small groups (typically < 20 individuals) in limited areas of dry deciduous forest, but across a vast territory from Sri Lanka to Myanmar. If current levels of forest protection are maintained, chance of extinction in the wild is not very high.

**Phenology:—**Leaf fall and flowers in December–March, followed by capsules.

**Etymology:—**The name was taken from Rheede (1679), adopting the “Brahmin” name “Nivuli” for the plant.

**Vernacular:—**Tamil and Malayalam: *Ela-kalli*; Kannada: *dubbakalli, dundukalli, elagalli*; Telegu: *akujemudu, akukall*; Hindi: *senhur, sij, thor*; Singalese: *kolapathok*; Burmese: *sha-soung*.

**Uses:—**It has medicinal uses like purgative, diuretic, poultice for fractures, wound healing, ear ache, etc.

**Notes:—**Although the plants were originally described from the Malabar region of Kerala in the 17^th^ century, we were unable to find such populations in Kerala today.

**Taxonomic notes:—**Hamilton separated out as a new species “Tithymalus indicus spinosus, nerii folio”, J. Commelin (1697: 25, plate 13) and “Ela calli” Rheede, (1689: 83, plate 43) from *E. neriifolia* L. on the basis that this plant has terete branches rather than 5-angular branches. Different authors, as listed above under misapplied names, used the name *E. neriifolia* incorrectly to describe what is actually *E. nivulia*.

Dorsey *et al*. (2013) included a plant labeled as *Euphorbia teke* Schweinf. ex Pax (1894: 118) in their genetic study of *Euphorbia* sect. *Euphorbia*. Even though this is an African taxon, the markers placed it well within the Indian clade. According to their electronic supplement, the sample they used came from a horticultural source at U.C. Davis in the US. Similarly, Zimmerman (2010), used plant material from a horticultural source in South Africa, showing *E. teke* to be closely related to *E. nivulia*. A paper on *E. teke* by Minch (1995) presents photos that do not match the herbaria specimens of the species and are clearly *E. nivulia.* It appears that all three sources have used samples of *E. nivulia* from India, incorrectly identifying them as *E. teke*. We are confident that actual material of *E. teke* from tropical Africa would fall well within the African clade.

***Euphorbia paschimia*** Malpure, Sardesai & B. DeJong, *sp. nov*.

TYPE**:—** INDIA. Maharashtra: Ahmednagar district, Chandanapuri Ghat, N 19°26’26.01“ E 074°12’03.97”, elev. 727 m, 18 February 2018, *N.V. Malpure 053* (holotype CAL!; isotypes BSI!, SUK!).

Misapplied names:

*Euphorbia neriifolia* auct. mult. *non* Linnaeus (1753: 451): Hooker (1887: 255); Singh (2002: 48); Eflora of India (2007: *Euphorbia neriifolia*); Binojkumar & Balakrishnan (2010: 310); Sankara Rao *et al*. (2019: *Euphorbia neriifolia*); Digital Flora of Gujarat (N.D.: *Euphorbia neriifolia*).

**Diagnosis:—**Densely crowded, spreading, flat-topped lactiferous shrub to small tree with medium sized leaves crowded near the ends of branches. Segments usually short with long, narrow, sub-confluent tubercles arranged in 5–7 distinctly spiral rows. New growth variegated. Leaves oblanceolate. Cymes with very short peduncles, frequently 3 secondary peduncles from a single primary peduncle, cyathia small, deep red, capsules triangular, seeds small and dark brown. *E. neriifolia* differs from it in having rounded crown, much smaller tubercles, spoon-like leaves, does not have variegated new growth, has larger rust colored glands, and does not usually produce capsules.

**Description:—**Densely crowded, spreading, flat-topped shrub 2–4 m, occas. a small tree up to 6 m with sparse rounded crown. *Bark* grey with persistent tubercle and branch scars. *Branches* vertically oriented with short segments, 5–20 cm, variegated new growth. *Tubercles* prominent, variable length, 5–15 mm, sub-confluent, narrow with flattened sides. *Spine shields* dark brown becoming dark grey, circular, 5–6 mm, with divaricate spines in a pair, 2–8 mm. *Leaves* succulent, crowded at the ends of branches, 6–8 × 2.5–3.5 cm, entire with edge sometimes wavy, oblanceolate, apex obtuse, apiculate, base cuneate tapering into petiole 0.5 cm, prominent central nerve, lateral nerves faint, deciduous during the dry season. Single *cyme* from flowering eye, 1-2 mm above spine shield, short peduncles 2–5 mm, often 3 (4) secondary peduncles per single primary, branching up to 4 times, *bracts* broadly crescent shaped, 2.5 mm, reddish with short lacerate fringe ending below gland, no keel, central *cyathium* staminate, lateral cyathia bisexual, *involucre* cupular, 4–5 × 4–5 mm, glands narrow oblong, deep red, 2.5 mm, lobes broadly oval, fimbriated edges, 2 × 2.5 mm. *Male flowers* in 5 fascicles of 8– 10 flowers, pedicel translucent 1.5 mm, filament greenish 0.5–0.7 mm,, anthers yellow sub-globose, 0.5 mm, fissure longitudinal, bracteoles narrow, lacerate,1.5 mm. *Female flower* pedicel approx. 1 mm, ovary subglobose with 3 lobes, 2 × 2 mm, styles split into 3 at base, bifid, stigma papillose, *capsule* excerted from the involucre, pedicel of capsule retroflexed when immature becoming curved or straight, approx. 10 mm, capsule, red becoming pinkish to yellowish, 10–12 mm, triangular, visible sutures between cocci, ends of cocci acute, *seeds* sub-globose, 2.5 mm, brown without spots. [Figs. 11, 12]

**FIGURE 11.**
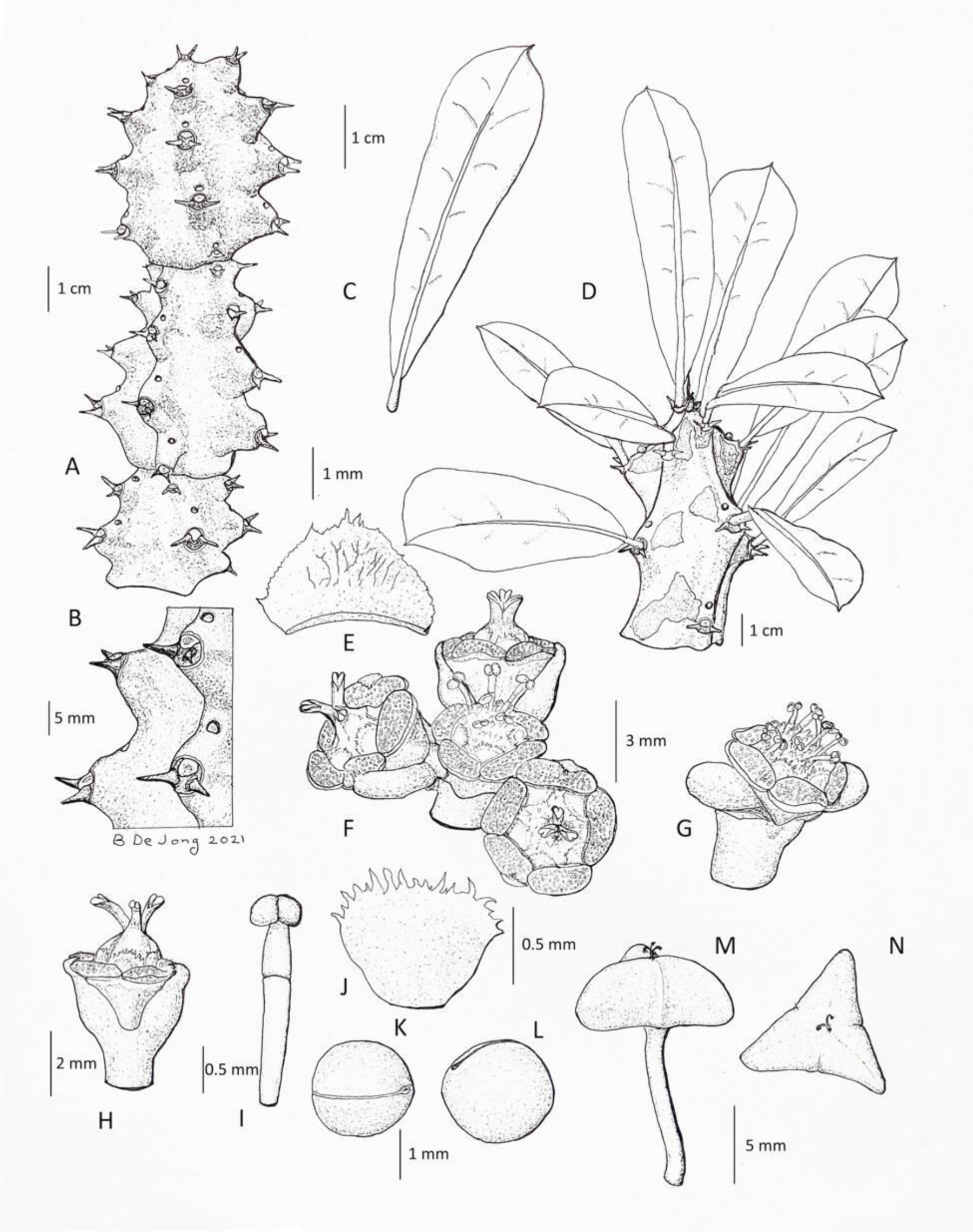
*Euphorbia paschimia* Malpure, Sardesai & B. DeJong (A) Habit, (B) Tooth and spine shield detail, (C) Leaf, (D) Growing branchlet, (E) Bract, (F) Cyme with triad of bisexual cyathia, (G) Staminate cyathium lateral, (H) Bisexual cyathium lateral, (I) Male flower, (J) Lobe of cyathium, (K) Seed raphe view, (J) Seed side view, (L) Capsule lateral view, (M) Capsule top view. Illustrated by Bruce De Jong from N. V. Malpure 053.

**FIGURE 12.**
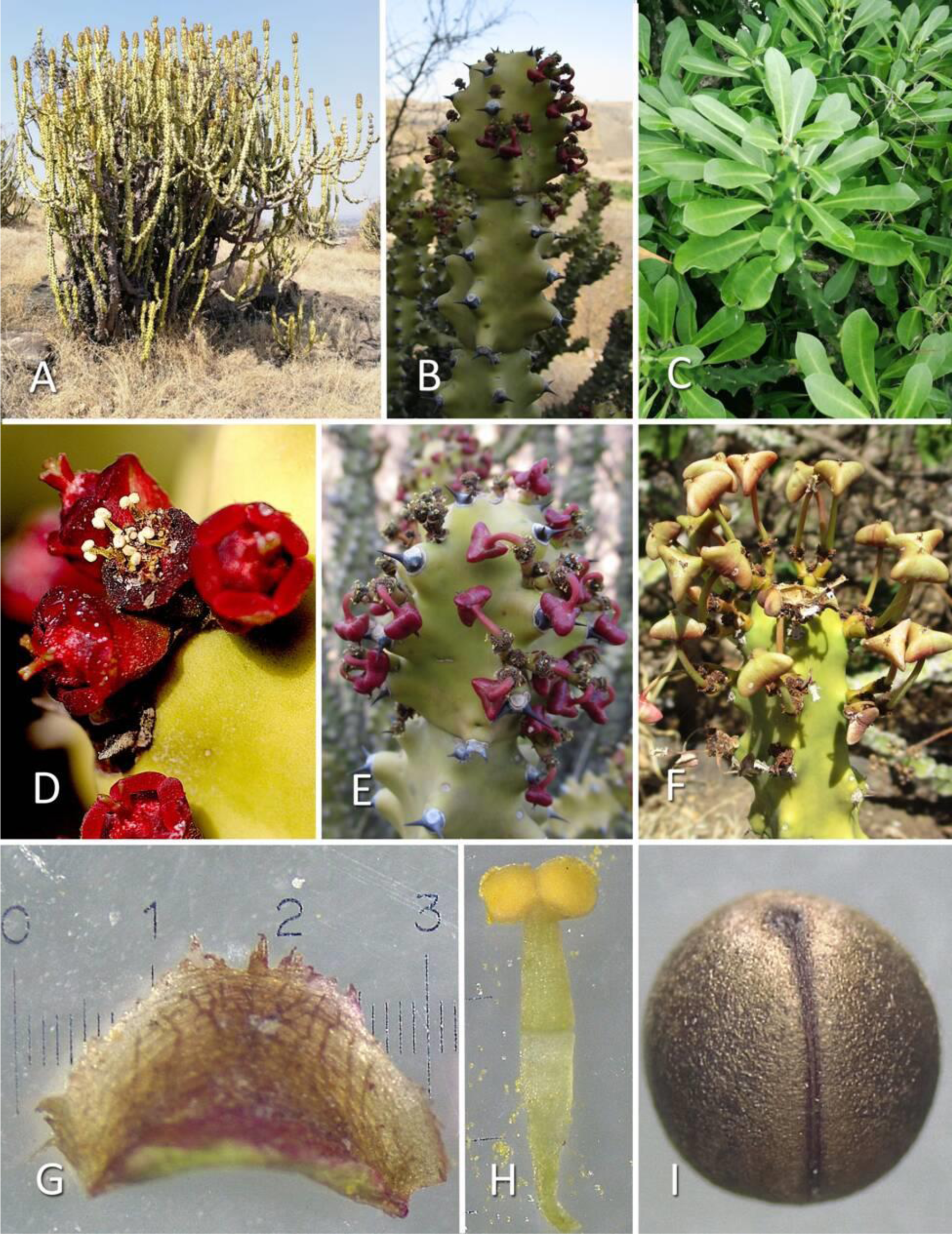
*Euphorbia paschimia* Malpure, Sardesai & B. DeJong, (A) Habit, (B) Branchlet, (C) Leaves and new growth showing variegation, (D) Cyme with triad of bisexual cyathia, (E) Early fruiting branch, (F) Mature capsules, (G) Bract, (H) Male flower, (I) Seed. Photos A–I by N. V. Malpure.

**Distribution:—**Western India: Maharashtra, Daman & Diu, Karnataka, Gujarat, Madhya Pradesh, and probably Rajasthan. [See Map 1.]

**Specimens examined:—** INDIA. Daman & Diu: Daman Dt., Varkunda village, 4 March 1963, *S.R. Rolla* 88967 (BSI 88967!); Gujarat: Bharuch Dt., Amod, 19 January 1894, *P.R. Mehta 1727* (CAL 397800!); Madhya Pradesh: Khargone Dt., on way to Kundi village 13 February 1987, *M. Prasad 39505* (BSA 9505!); Tikamgarh Dt., Baldeogarh village, 19 April 1989, *G.P. Roy & K. Kishore 44773* (CAL0000008937!); Maharashtra: Bhandara Dt., Way to Dakhni, 11.12.1977, *S. K. Malhotra 151279* (BSI 100874!); Pimpalgaon, 9 March 1983, *P.G. Diwakar* 164991 (BSI 164991!); Dhule Dt., Sita khai surrounding, 9 May 1964, *K.K. Ahuja 96753* (BSI 96753!); Nandurbar Dt., Ashte village, 5 January 1957, *S.K. Jain 11060* (BSI 15425!); Nashik Dt., 56^th^ Mile Point Mumbai to Taloda road, 6 May1964, *S.R. Rolla 96649* (BSI 73574!); Trimbak, 3.2.1983, *R.L. Narasimhan* 165306 (BSI!); Pune Dt., Ambavane, 31 March 1964, *B.V. Reddi 97693* (BSI 97693!); Khandala Hills, 15 March 1959, *S.K. Mukherjee 52243* (BSI 15422!); Sindhudurg Dt., Amboli west, 17 November 1902, *collector unknown 17070* (CAL 397803!); Kankavli taluk, Sangave village, 14 April 1971, *B.G. Kulkarni 128716* (BSI 100279!); Solapur Dt., Barshi to Ramling forest, 21.7.1957, *S.K. Jain 20108* (BSI 20108!); Thane Dt., Savdhani Hill, Mandir Range, 18 January 1968, *K.V. Billore 113636* (113636!); Murbad village, Daheri range, shrubs growing along the borders of cultivated fields 13 April 1968, *K.V. Billore 113920* (BSI 113920!).

**Habitat:—**Dry rocky hills of the western Deccan plateau, growing in scrub jungle and secondary forest. Also extends to the wetter coastal ranges of the Western Ghats, from near sea level up to 1500 m.

**Conservation status:—**Considering the broad range of the plants, across at least 1000 km north to south and east to west, our subjective impression is that the species would be Least Concerned. This could change with continued habitat threat.

**Phenology:—** Leaf fall and flowers in December–March, followed by capsules.

**Etymology:—**Specific epithet is the vernacular word “paschim” for west, based on the location in western India.

**Vernacular:—**Marathi: *Sabar, Sabarkand*

**Uses:—** It is used as a purgative, diuretic, poultice for fractures, wound healing, for removing warts, for ear ache, and as an insecticide (Binojkumar & Balakrishnan 2010).

**Notes:—** This plant is polymorphous. In full sun on rocky hills, it tends to have short thick, tightly spiraled segments and grows as a spreading shrub. In wetter environments or in partial shade, fence lines, or in competition with taller plants, it can have long, thin, winding segments and can grow into a sprawling tangled shrub, or become small tree up to 6 meters. Its main distinguishing features, however, are retained.

***Euphorbia yadavii*** Malpure, Raut & B. DeJong, *sp. nov*.

TYPE:**—** INDIA. Maharashtra: Ahmednagar district, Rampurwadi, N 19 43’ 9.5808’’ E 74 35’ 6.1548’’, elev. 513 m, 29 April 2018, *N.V. Malpure 046* (holotype CAL!; isotypes BSI!, SUK!).

Misapplied names:

*Euphorbia neriifolia* auct.*mult. non* Linnaeus (1753: 451): Beddome (1872: 216); Talbot (1902: 207); Brandis (1906: 558); Burkill (1910:130); Stewart (1874: 439); Gamble (1925: 1277); Parker (1918: 446); Haines (1921: 242); Radcliffe-Smith (1986: 172); Rao & Sreeramulu (1986:423); Bhat (2003: 564).

**Diagnosis:—**Shrub to small tree allied to *E. nivulia,* but differing in its irregular broadly columnar crown, thick vertically oriented medium length segments, prominent rounded tubercles arranged in 5 strongly spiral rows, spine shields large and oval, with long spines, cyathia light yellow on medium length peduncles, glands elliptical, capsules yellowish to pinkish, deeply lobed with less prominent sutures between cocci, seeds dark grey with light grey spots.

**Description:—**Small lactiferous, succulent tree 4–6 m, irregular broadly columnar crown, highly whorled. *Bark* light grey-brown, rugose. *Branches* medium length 20–50 cm, stout with marked narrowing towards base, prominent tubercles 5–8 mm, conical or slightly flattened sides, arranged in 5 strongly spiral rows. *Spine shields* brown, oval, 5 × 8 mm, with divaricate spines in a pair up to 8 mm. *Leaves* succulent, crowded at the ends of branches, 8–24 × 4–6.5 cm, flat to slightly spoon-like, entire, obovate with rounded or obtuse apex, apiculate, base cuneate tapering to 1 cm petiole, prominent central nerve, indistinct lateral nerves, deciduous during the dry season. Single *cyme* per flowering eye, 1–2mm above spine shield, primary peduncle 9–15 mm, secondary and tertiary peduncles 10–12 mm, *bracts* 3–4.5 × 4–5 mm, oval, greenish broadly ovate 3 × 3 mm, not keeled, central *cyathium* staminate, lateral cyathia bisexual, usually one branching but up to two in mature trees, involucre shallow cupular, 10–12 × 4–5 mm, *glands* somewhat elliptical, flattened medially, light yellow, 3.5–5 mm, *lobes* broadly ovate, 1.5–2 mm, sl. lacerate edges. *Male flowers* in 5 fascicles of 6–7 flowers, pedicel translucent 4 mm, filament pinkish 1 mm, anthers pink in a pair, reniform, 0.7 mm, fissure longitudinal, bracteoles narrow spathulate, highly lacerate, 1.5– 2.0 mm. *Female flower* pedicel approx. 2 mm, ovary somewhat broad with 3 lobes, 2 × 3 mm, styles connate to middle, widened at tips, stigmata papillose, *capsule* exerted from the involucre, pedicel of capsule retroflexed when immature becoming curved or straight, approx. 15–18 mm, capsule light green to pinkish, 15 × 5 mm, deeply lobed, suture between cocci not prominent, ends of cocci skewed downwards, *seeds* globose, 3.0 mm, dark grey with light grey spots coalescing into the raphe [Figs. 13, 14].

**FIGURE 13.**
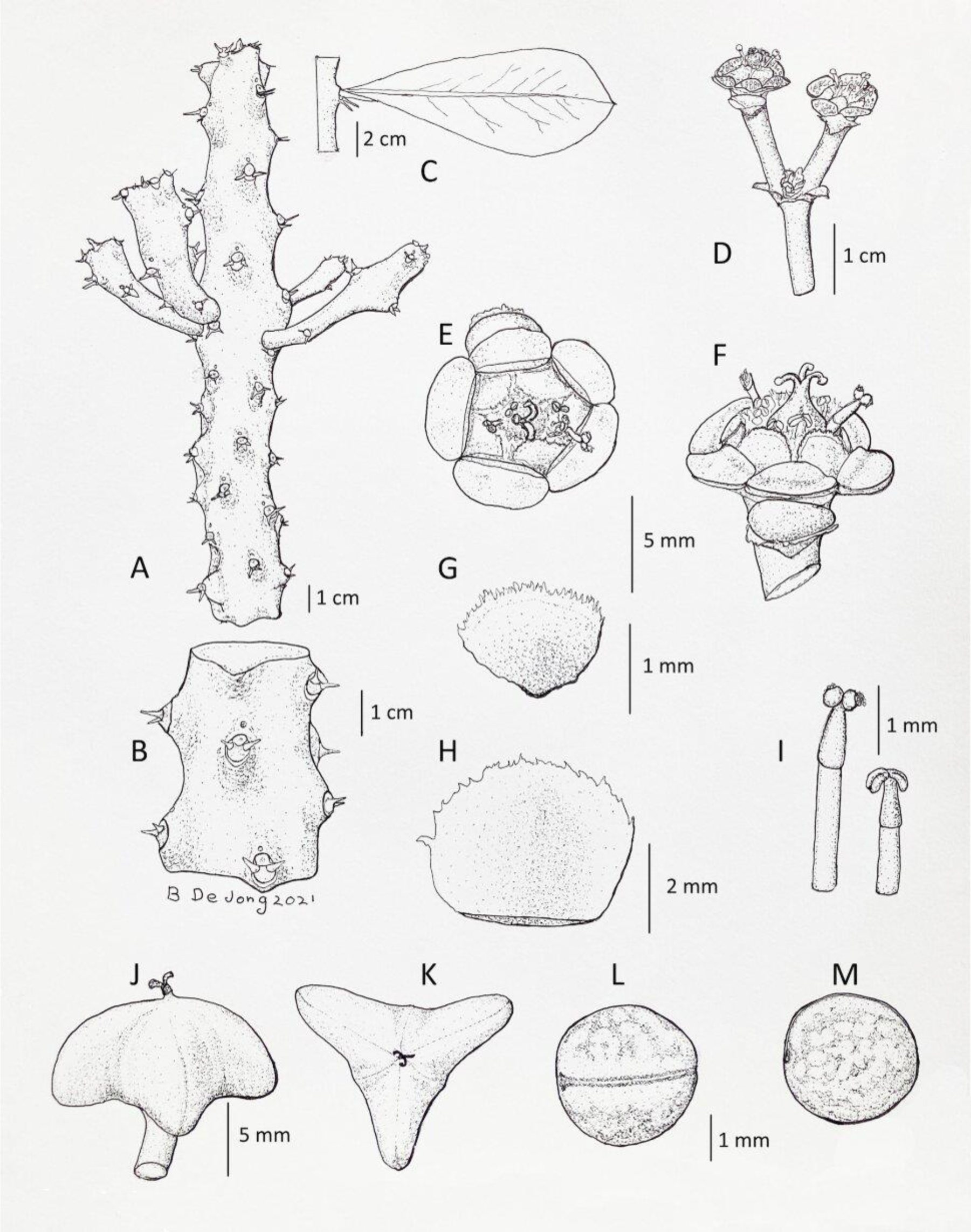
Illustration of *Euphorbia yadavii* Malpure, Raut, & B. DeJong (A) Habit, (B) Tubercles and spine shields close-up, (C) Leaf, (D) Simple cyme, (E) Bisexual cyathium top, (F) Bisexual cyathium lateral, (G) Lobe of cyathium, (H) Bract, (I) Male flowers, (J) Capsule lateral, (K) Capsule top, (L) Seed raphe view, (M) Seed side view. Illustration by Bruce De Jong from N. V. Malpure 046.

**FIGURE 14.**
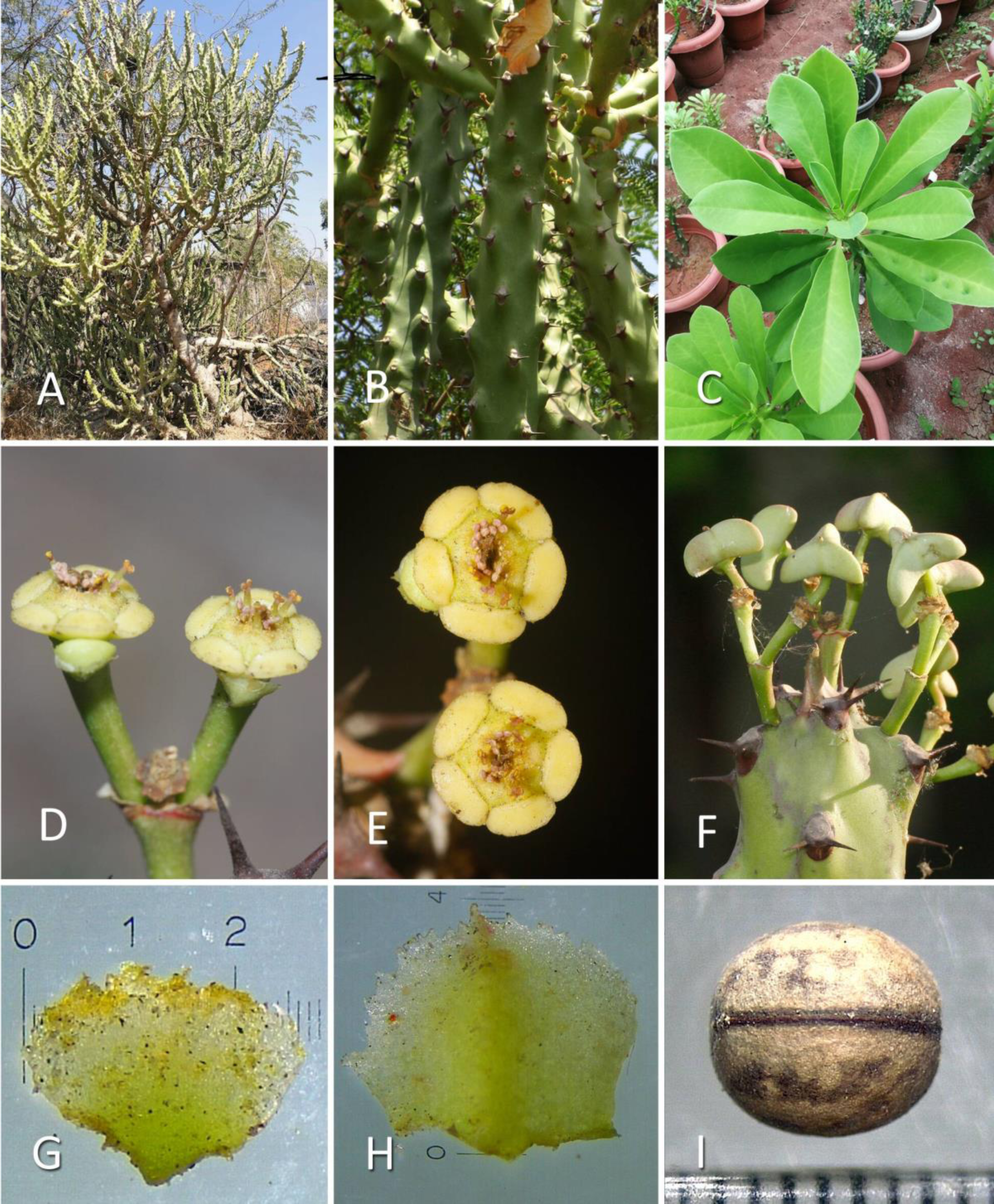
*Euphorbia yadavii* Malpure, Raut & B. DeJong (A) Habit, (B) Branchlets close-up, (C) Leaves, (D) Cyme, (E) Bisexual cyathia top view, (F) Fruiting branch, (G) Lobe of cyathium, (H) Bract, (I) Seed raphe view. Photos A–I by N. V. Malpure.

**Distribution:—**Western India, especially Maharashtra, but also recorded in Haryana, Rajasthan, Gujarat, and Madhya Pradesh. Mostly seen in cultivation as fence-lines. Wild origin obscure, possibly Rajasthan. [See Map 1.]

**Additional specimens examined:—**INDIA. Dadra & Nagar Haveli Dt.: Dhapsa, 16.11.1970, *M.Y. Ansari 127198* (BSI 87855!) Gujarat: Ahmedabad, October 1899, *Kalkaprasav 12694,* (CAL 397794!); Haryana: Rohtak Dt., Rohtak, 18 January 1903, *J.H. Burkill 18172* (CAL 397810!); Madhya Pradesh: Khargone Dt., on the way to Dharampuri, 10 February 1987, *M. Prasad 39455* (CAL 000008945!); Bhilwara Dt., Kasthurbagram Krishi Kshethra, near guest home, 30 January 1987, *K.K. Khanna & R. Saran 38971*, (BSI 38971!); Maharashtra: Akola Dt., Patur Taluk, Medshi village, 21.2.1971, *S.Y. Kamble 15271*1 (BSI 106033!); Yavatmal Dt., Yavatmal, 29.8.1907, *C.A. Malcolm 27545* (CAL 379829!); Rajasthan: Bhilwara Dt., Meja Dam, 15 February 1979, *A.N. Singh 7136*, (CAL 0000008944!)

**Habitat:—**Currently found across cultivated areas on the plains of western India.

**Conservation status:—** Our subjective impression is that with the ease of cultivation and wide-spread use in fence-lines, it is not vulnerable.

**Phenology:—**Leaf fall and flowers in December–March, followed by capsules.

**Etymology:—**The name is in honour of prominent Indian botanist S. R. Yadav.

**Vernacular:—** Marathi: *Sabar, Sabarkand*

**Uses:—** Fence-lines

**Taxonomic notes:—**This plant, by virtue of having 5 rows of spiral tubercules and leaves bunched at the end is usually identified as *E. neriifolia.* It differs however in its geography, habit, leaf morphology, tubercles that are not confluent, large oval spine shields, large spines, cymes with long peduncles only branching up to 2 times, yellow cyathia with elliptical glands, and the presence of capsules and seeds. Its nearest relative botanically appears to be *E. nivulia,* but this species is a larger tree with a large rounded crown, heavily fissured corky bark, spreading straight terete branches, longer peduncles, greenish cyathia with red filaments and dark red anthers, and larger brown spotted seeds. *E. yadavii* is easily differentiated from *E. paschimia* which is also frequently identified as *E. neriifolia*. (See the comparison table).

## Conclusions

After having analysed the world literature on the *E. neriifolia* complex and conducting field studies in India, Indonesia, and Africa, we have limited *E. neriifolia* to the type plants originally cultivated in China and S E Asia. Proposed an epitype to stabilise *E. neriifolia,* reinstated *E. ligularia* as a species and synonymized *E. undulatifolia* under *E. ligularia.* Designated a neotype for *E. drupifera* and a neotype for *E. complanata* which we have synonymised under *E. neriifolia.* Furthermore, separated out and described two new species *E. yadavii* and *E. paschimia*from India which have been very commonly confused with *E. neriifolia.*:

### Key to the *E. neriifolia* complex and related species

1. Leaves < 6 cm … [rest of South Asian *Euphorbia* section *Euphorbia*] – Leaves > 6 cm … 2.
2. Spine shields on tubercles … 3.

– No spine shields or tubercles, presence of underground tubers … [Indian geophytes]
3. Tubercles arranged on vertical wings … 4

– Tubercles arranged in lines, not on wings… 5
4. Wings, 5–7, flat between wings, leaves not persistent, fruit a capsule … *E. royleana*

– Wings (4) 5 (–6), not flat between wings, leaves persistent, fruit a dupe … *E. drupifera*
5. Tubercles in spirals, sub-confluent … 6

– Tubercles in spirals, not sub-confluent … 7
6. Tubercles, 1–4 mm in 5 rows, leaves spoon-like concave upwards, glands reddish, tree with rounded crown … *E. neriifolia*

– Tubercles 5–8 mm in 5 rows, leaves flat and sl. drooping, glands yellow-green, tree with columnar crown… *E. ligularia*
– Tubercles 5–15 mm in 5–7 rows, leaves flat to sl. concave upwards, cyathia small, glands deep red, spreading flat-topped shrub … *E. paschimia*
7. Spine shields circular … 8

– Spine shields not circular … 9
8. Tubercles 1–3 mm long, branchlets terete, sp. shields with one pair of spines, glands greenish yellow … *E. nivulia*

– Tubercles up to 10 mm long, sp. shields with 2 pairs of spines, leaves broadly elliptical with prominent veins … *E. sahyadrica*
9. Tubercles 5–8 mm long, sp. shields oval, glands light yellow … *E. yadavii*

– Tubercles 10 mm, sp. shield broadly horizontal/trapezoidal, split into two, cyme branching up to 4 times, glands reddish, pygmy habit … *E. lakshminarasimhanii*

## Acknowledgements

The authors (NVM & PSR) are grateful to the Science and Engineering Research Board (SERB), New Delhi (EMR/2016/001574) for providing financial assistance and to the Principal, SSGM College, for encouragement and providing all the necessary facilities. MMS is thankful to University Grants Commission, New Delhi for funding under the University with Potential for Excellence (UPE) program and to the authorities of Savitribai Phule Pune University, Pune. We also thank the curators of the following herbaria: BAMU, CAL, MH, DD, BSI, SUK, JCB, and RHT as well as the editor and reviewers for their critical comments and revision of the manuscript.

In addition, we would like to thank the following for permission to reproduce images: Allard Pierson Depot–University of Amsterdam, The Linnaean Society, Digital Images–The Real Jardín Botánico Spain, the Curator Images–Royal Botanical Gardens Kew, the Missouri Botanical Garden Press, and the Science Press–Beijing, and the Collections Manager Naturalis Biodiversity Center Holland. We are grateful to Geoff Stein for his photos, to Mr. Geda of Bali Indonesia for assistance in finding *E. neriifolia* in the field, to Dr. Sunil Kaul for assistance in finding *E. ligularia* in Bodoland Assam, to the Iora Hotel, Kasziranga Assam for providing samples of *E. ligularia,* to Olaf Ryding, Collection Manager, Danish Herbarium for information on type materials of *E. drupifera*, to Nicolien Sol and Peter van Wiesen of the Naturalis Biodiversity Center for information on type materials of *E. undulatifolia*, to Jörn Hentschel of Herbarium JE and Robert Vogt of Herbarium B for assistance with specimens of *E. camplanata*, and to Elizabeth Charmentier and Mark Spencer of the Linnaean Herbarium for information on Osbeck’s Linnaean specimens. We also thank Dr. K.N. Gandhi Department of Organismic and Evolutionary Biology Harvard and Dr. John McNeill of Royal Botanic Garden of Edinburgh for their taxonomic advice.

## References

Ainslie, W. (1826) Materia Medica 2. Longman, Rees, Orme, Brown, & Green, London, 604 pp.

Aiton, W. (1789) Hortus Kewensis, Vol. 2. George Nicol, London, 460 pp.

Aubriet, C. (1690 – 1735) Collections de Velins du Museum National d’Histoire Naturelle, Vol. 60, t. 28. Available from: http://www.plantillustrations.org/illustration.php?id_illustration=280119&uhd=2&mobile=0 (accessed: 10 November 2022).

Backer, C.A. & Bakhuizen van den Brink, R.C. (1963) Flora of Java, Vol. 1. N.V.P. Nordhoff, Groningen, The Netherlands, 668 pp.

Balakrishnan, N.P. & Chakrabarty, T. (2007) *The family Euphorbiaceae in India – a synopsis of its profile, taxonomy and bibliography*. Bishen Singh Mahendra Pal Singh, Dehra Dun, India, 500 pp.

Bamber, C.J. (1916) Plants of the Punjab. Superintendent, Government Printing, Lahore, 652 pp.

Bedomme, R.H. (1872) The Flora Sylvatica for Southern India, Vol. 3. Ganz Brothers, Madras, India, 319 pp.

Berger, A. (1907) Sukkulente Euphorbien. Eugen Ulmer, Stuttgart, 132 pp.

Bhat, K.G. (2003) Flora of Udipi District. Poorna Prajna College Publications, Udipi, Karnataka, India, 913 pp.

Binojkumar, M.S. & Balakrishnan, N.P. (2010) The Genus Euphorbia L. (Euphorbiaceae) in India, A Taxonomic Revision. Bishen Singh, Mahendra Pal Singh, Dehra Dun, India, 430 pp.

Blanco, M.F. (1837) Flora de Filipinas. In the press of St. Thomas, for D. Candido Lopez, Manila, 886 pp.

Blatter, E. & Hallberg, F. (1919) The Flora of the Indian Desert (Jodhpur and Jaisalmer). Journal of the Bombay Natural History Society 26: 970.

Boerhaave, H. (1727) Index Plantarum. Janssonios Vander Aa, Leiden, 320 pp

Boissier, P.E. (1862) Sistens Euphorbiaceas. *In*: De Candolle, A. (Ed.), Prodromus Systematis Naturalis Regni Vegetabilis, Vol. 15 (2). Victoris Masson et Fillii, Paris, 1286 pp.

Bonelli, G. (1772) Hortus Romanus, Vol. 1, t. 28. Bouchard and Gravier, Rome, 100 pp.

Bradley, R. (1725) The History of Succulent Plants, Decade 111. William Mears, London, 135 pp.

Bretschneider, E. (1880) Early Researches into the Flora of China. Journal of the North-China Section of the Royal Asiatic Society, 1: 90.

Breyn, J. (1739) Prodromii Fasciculi Rariorum Plantarum. David Fridericus Rhedius, Gedansk, 108 pp.

Brown, N.E. (1913) Euphorbiaceae. *In*: Thiselton-Dyer, W.T. (Ed.), Flora of Tropical Africa, Vol. 6 (1). Lovell Reeve and Co., London, 1094 pp.

Bruyns, P.V., Mapaya, R.J. & Hedderson, T. (2006) A new subgeneric classification for *Euphorbia* (Euphorbiaceae) in Southern Africa based on ITS and *psbA-trnH* sequence data. Taxon 55: 397–420. 10.2307/25065587

Buchanan-Hamilton, F. (1825) Dr. Francis Hamilton’s commentary. Transactions of the Linnean Society of London Vol 14 (2). Richard Taylor, London, 605 pp.

Burkhill, I.H. (1910) Notes from a Journey to Nepal. In: Records of the Botanical Survey of India, Vol. 4 (4). Botanical Survey of India, Calcutta, 140 pp.

Burman, J. (1737) Thesaurus Zylanicus. Janssonio-Waesbergios & Salomonem Schouten, Amsterdam, 235 pp.

Burman, N.L. (1768) Flora Indica. Johann Schreuder, Amsterdam, 260 pp.

Carter, S. (2002) *Elaeophorbia*. In: Eggli, U (Ed.). Illustrated Handbook of Succulent Plants: Dicotyledons. Springer-Verlag, Berlin, 547 pp.

Chevalier, A. (1948) Plants crassulascents de l’Ouest africain (Opuntias et Euphorbes). Journal d’agriculture traditionnelle et de botanique appliquée, Bulletin No. 309–310: 347–348.

Commelin, J. (1697) Horti Medici Amstelodamensis. P. & J. Blaeu and Abrahamum Somerem, Amsterdam, 220 pp.

Commelin, C. (1703) Praeludia Botanica. Fredericum Haringh, Leiden, 129 pp.

Cooke, T. (1908) Flora of the Presidency of Bombay, Vol. 2. Taylor and Francis, London, 1083 pp.

Croizat, L. (1934) De Euphorbio Antiquorum Atque Officianarum. Levi, Simpson & Company, New York, 127 pp.

DeFilipps, R.A. & Krupnick, G.A. (2018) The medicinal plants of Myanmar. PhytoKeys 102: 114.

Dorsey, B.L., Haevermans, T., Aubriot, X., Morawetz, J.J., Riina, R., Steinman, V.W. & Berry, P.E. (2013) Phylogenetics, morphological evolution, and classification of *Euphorbia* subgenus *Euphorbia*. Taxon 62(2): 291–315.

Burger, W. & Huft, M. (1995) Family # 113 Euphorbiaceae. In: Burger, W. (Ed.) Fieldiana Botany New Series no. 36, Flora Costaricensis. Field Museum of Natural History, Chicago, 169 pp.

Digital Flora of Gujarat State. (N.D.). Euphorbia nivulia. Available from: http://gujaratflora.com/showplantdescription1.php?plantname=%3Cp%3E%3Cem%3EEuphorbia%20nivulia%3C/em%3E%20Buch.-Ham.%3C/p%3E (accessed: 3 May 2020).

Duthie, J. F. (1915) Flora of the Upper Gangetic Plain. Superintendent Government Printing, Calcutta, 283 pp.

Efloraofindia (2007 onwards) Euphorbia neriifolia. Available from: https://sites.google.com/site/efloraofindia/species/a---l/e/euphorbiaceae/euphorbia/euphorbia-neriifolia (accessed: 3 May 2020).

Ekanayake, S. P., Pieris, T. N., Jayatilake, W., Goonatilake, S., Asela, M.D.C., Pieris, A. L., & Bandara, A. (2015) *Euphorbia Nivulia* Buch.-Ham. (Euphorbiaceae) – a new addition to the flora of Sri Lanka. Zoo’s Print 30(6): 30–31.

Esser, H. –J. (2005) 37. *Euphorbia*. *In*: Chayamarit, K. & Welzen, P. C. van, Euphorbiacae. *In*: Santisuk, T. & Larsen, K. (eds.) Flora of Thailand, Vol. 8(1), pp. 290–291. The Forest Herbarium. Bangkok, 303 pp.

Esser, H. –J. (2017) Euphorbia. *In*: Welzen, P. V. van, Flora of Thailand, Euphorbiaceae. Available from: http://www.nationaalherbarium.nl/ThaiEuph/ThEspecies/ThEuphorbiaT.htm (Accessed: 8 May 2020).

Forman, L.L. (1997) Notes concerning the typification of names of William Roxburgh’s species of phanerograms. Kew Bulletin 52: 513–534.

Gagnepain, F. (1910) Euphorbiacées. *In*: LeCompte, M. H. (Ed.), Flore Générale de L’Indochine, Vol. 5, Masson et Cie, Paris, 324 pp.

Gamble, J.S. (1915) Flora of the Presidency of Madras, Vol. 3. Adlard and Son, London, 2017 pp.

Haines, H.H. (1925) Flora of Bihar and Orissa. Adlard & Son & West Newman, Ltd. London, 1350 pp.

Hansen, C. & Maule, E.F. (1973) Per Osbeck’s collections and Linnaeus’s *Species Plantarum* (1753). Botanical Journal of the Linnaean Society 67: 189–212.

Haworth, A.H. (1812) Synopsis Plantarum Succulentarum. Richard Taylor and Friends, London, 207 pp.

Haworth, A.H. (1828) Description of new succulent plants. In: Tailor, R. & Phillips, R. (Eds.) The Philosophical magazine or annals of chemistry, mathematics, astronomy, natural history, and general science, pp. 183–188.

He, D. (1997) Euphorbia neriifolia, Fig. 352. Available from: http://www.efloras.org/object_page.aspx?object_id=109316&flora_id=2 (Accessed: 1 April 2020).

Hermann, P. (1698) Paradisus Batavus. Abrahamum Elzevier, Leiden, 247 pp.

Hermann, P. (1717) Museum Zylanicum. Isaacum Severinum, Leiden, 71 pp.

Ho, P. H. (1992) Cayco Vietnam, an illustrated flora of Vietnam, vol. 2, p. 358, fig. 4663. Pham-Hoang Ho, Montreal, 951 pp.

Hoeck, F. (1896) Allgemeine Pflanzengeographie. *In*: Koehne, E., Just’s Botanischer Jahresbericht, Vol. 11. Gebrueder Borntraeger, Berlin, 650 pp.

Hooker, J.D. (1890) The Flora of British India, Vol. 5. L. Reeve and Co., London, 910 pp.

Hooker, J.D. (1898) Handbook to the Flora of Ceylon, Vol. 4. Dulau and Co., London, 384 pp.

Houtyn, M. (1777) Natuurlyke Historie of Uitvoerige Beschriving der Dieren, Planten, en Mineraalen, Vol. 2, Part 8. F. Houtyen, Amsterdam, 784 pp.

Hutchinson, J. & Dalziel, J.M. (1952) Flora of West Tropical Africa, second edition,Vol. 1. Crown Agents for Overseas Governments and Administrations, London, 295 pp.

Isnard, (1720) Botanique. In: Collection Academique etc. Vol. 5. Hotel de Thou, Paris, 498 pp.

IUCN Standards and Petitions Committee (2022) Guidelines for Using the IUCN Red List Categories and Criteria. Version 15.1. Prepared by the Standards and Petitions Committee. Available from: https://www.iucnredlist.org/resources/redlistguidelines, (accessed 10 November 2022).

Janse, J.A. (1953) *Euphorbia undulatifolia* Janse spec. *nov*. The National Cactus and Succulent Journal 8(4): 69–70.

Jarvis, C.E., (2007) Order Out of Chaos: Linnaean Plant Names and Their Types. Linnaean Society of London, London, 1016 pp.

JSTOR (2013) Buchanan-Hamilton, Francis (1762–1829). Available from: https://plants.jstor.org/stable/history/10.5555/al.ap.person.bm000324521 (accessed 21 November 2022).

Kanjilal, U.N, Kanjilal, P.C., De, R.N., & Das, A. (1940) Flora of Assam, Vol. 4. R. C. Roy Chaudhurry, Calcutta, 395 pp.

Karsten, H. (1882) *Deutche Flora*. Pharmaceutisch-medicinische Botanik. J.M. Spaeth. Berlin, 1284 pp.

Kiggelaer, F. (1690) Horti Beaumontiani Exoticarum Plantarum Catalogus. The Hague, 42 pp.

Kurz, S. (1877) Forest Flora of British Burmah, Vol. 2. Superintendent of Government Printing. Calcutta, 613 pp.

Lamaire, C.H. (1786) L’illustration Horticole des Serres et des Jardin, Vol. 4. F.& E. Gyselinck, Ghent, 110 pp.

Lamarck, J.-B. (1786) Encyclopedie Methodique Botanique, Vol. 2. Panckoucke, Paris, 774 pp.

Linnaeus, C. (1737) Hortus Cliffortianus. George Clifford, Amsterdam, 501 pp.

Linnaeus, C. (1747) Flora Zylanica. Laurentii Salvii, Stockholm, 240 pp.

Linnaeus, C. (1748) Hortus Upsaliensis. Laurentii Salvii, Stockholm 306 pp.

Linnaeus, C. (1753) Species Plantarum. Laurentii Salvii, Stockholm, 1200 pp.

Linnaeus, C. (1754) Herbarium Amboinense. L.M. Hojer, Upsala, 28 pp.

Loureiro, J. (1790) Flora Cochinchinensis, Vol. 1. Jussu Acad. Reg. Scient., Lisbon, 353 pp.

Ma, J., & Gilbert, M.G. (2008) 74. Euphorbiacae. *In*: Wu, Z. & Raven, P. (Eds.) Flora of China Vol. 11. Missouri Botanical Garden Press, St. Louis, pp. 288–313.

Malaysia Biodiversity Information System (MyBIS) (2015 onwards) Euphorbia Neriifolia. Available from: https://www.mybis.gov.my/sp/25417 (accessed 14 January 2020).

Malpure, N.V., Chandore, A.N. & Yadav, S.R. (2016) *Euphorbia gokakensis* (Euphorbiaceae) from sandstone formations in Karnataka, India. Nordic Journal of Botany 34: 380–383.

Malpure, N.V., Raut, P.S., Chandore, A.N. & De Jong, B.E. (2021a) *Euphorbia lakshminarasimhanii*: a new pygmy succulent species from Konkan region of Maharashtra, India. Nordic Journal of Botany 39(7): e03142. 10.1111/njb.03142

Malpure, N.V., Raut, P.S., Sardesai, M.M. & De Jong, B.E. (2021b) *Euphorbia sahyadrica* (Euphorbiaceae), a new species of succulent shrub from the wet zone of the northern Western Ghats, Maharashtra, India. Phytotaxa 500(4): 285–293. 10.11646/phytotaxa.500.4.4

Mane, R.N., Nerlekar, A.N., Lapalikar, S.A., Kambale, S.S. & Yadav, S.R. (2017) The taxonomic status of two geophytic *Euphoriba* species (Euphorbiaceae) from Maharashtra, India. Phytotaxa 307: 141–146.

Manjunatha, B.K., Krishna, V. & Pulliah, T. (2004) Flora of Davangere District, Karnataka, India. Regency Publications, New Delhi, 464 pp.

Matthew, K.M. (1983) The Flora of the Tamilnadu Carnatic, Vol. 2. The Rapinat Herbarium, Tiruchirapalli, India, pp. 651–1540.

Merrill, E.D. (1918) Species Blancoanae. Bureau of Printing, Manila, 423 pp.

Merrill, E.D. (1923) An Enumeration of Philippine Flowering Plants, Vol. 2. Bureau of Printing, Manila, 530 pp.

Merrill, E.D. (1935) A Commentary on Loureiro’s “Flora Cochinchinensis”. Transactions of the American Philosophical Society, 24(*2*),444 pp.

Miller, P. (1763) The Gardener’s Dictionary, 8th edition, Euphorbia No. 5, Phillip Miller, London, A–Z pp.

Minch, L.W. (1995) *Euphorbia teke* Schweinf.: a rare species from Zaire. The Euphorbia Journal 3: 95–98.

Miquel, F.A.G. (1859) Flora van Nederlansch Indie, Vol. 1. C.G. van der Post, Amsterdam, 1116 pp.

Miquel, F.A.G. (1860). Flora Indiae Batavae, Supplementum Primum, Sumatra, p. 183. C.G. van der Post, Amsterdam, 656 pp.

Moninckx, J. (1702) Moninckx Atlas, Vol. 5, t. 4 Available from: https://www.uvaerfgoed.nl/beeldbank/nl/xview/?identifier=hdl:11245/3.27221;metadata=moninckx+5#page/5 (accessed: 1 May 2020).

Moninckx, J. (1690) Moninckx Atlas, Vol. 5, t. 45. Available from: https://www.uvaerfgoed.nl/beeldbank/nl/xview/?identifier=hdl:11245/3.27221;metadata=moninckx+5#page/46 (accessed: 1 May 2020).

Moon, A. (1824) A Catalogue of the Indigenous and Exotic Plants Growing in Ceylon. Wesleyan Mission Press, Colombo, 196 pp.

Morison, R. (1699) Plantarum Historiae Universalis Oxoniensis. Theatro Sheldoniano, Oxford, 617 pp.

Osbeck, P. (1757) Dagbok Ofwer en Ostindsk Resa. Lor. Ludv. Grefing, Stockholm, 376 pp.

Osbeck, P. (1771) A Voyage to China and the East Indies. Vol 1. Benjamin White, London, 356 pp.

Parker, R. N. (1918) A forest Flora for the Punjab with Hazara and Delhi. Superintendent, Government Printing, Lahore, 577 pp.

Parker, R. N. (1927) Forest Flora for the Chakrata, Dehra Dun, and Saharanpur Forest Divisions. Government of India, Central Publications Branch, Calcuta, 558 pp.

Pax, F. (1894) Euphorbiaceae africane. In: Engler, A. (Ed.) Botanische Jahrbücher für Systematik, Pflanzengeschichte un Pflanzengeographie 18: 76–127.

Poisson, M. J. & Pax, M. J. (1902) Bulletin Museum D’Histoire Naturelle. Imprimerie Nationale, Paris, 659 pp.

Pelser, P.B., Barcelona, J.F. & Nickrent, D.L. (2011 onwards) Co’s Digital Flora of the Philippines. Available from: https://www.philippineplants.org/Families/Euphorbiaceae.html (accessed 14 January 2020.

Perry, L.M. (1980) Medicinal Plants of East and Southeast Asia. The MIT Press, Cambridge, Massachusetts, 620 pp.

Philcox, D. (1997) 29. *Euphorbia*. In: Dassanayake, M. (Ed.), A Revised Handbook to the Flora of Ceylon, Vol. 11. A. A. Balkema, Rotterdam, 420 pp.

Plukenet, L. (1696) Almagestum Botanicum. Plukenet, London, 420 pp.

Prain, D. (1903) Bengal Plants, Vol. 2. Botanical Survey of India, Calcutta, 670 pp.

Radcliffe-Smith, A. (1972) In: Airy Shaw, H.K. (Ed.), The Euphorbiaceae of Siam. Kew Bull. 26 (2): 191–363.

Radcliffe-Smith, A. (1980) *In*: Airy Shaw, H.K., The Euphorbiaceae of New Guinea. Kew Bull. Additional Series VIII: 1–231.

Radcliffe-Smith, A. (1981) *In*: Airy Shaw, H.K., The Euphorbiaceae of Sumatra. Kew Bull. 36(2): 239–374.

Radcliffe-Smith, A. (1982a) *In*: Airy Shaw, H.K., The Euphorbiaceae of Central Malesia (Celebes, Moluccas, Lesser Sunda Is.). Kew Bull. 37(1): 1–40.

Radcliffe-Smith, A. (1982b) Euphorbiacées. *In*: Bosser, J. (Ed.), Flore Des Marcareignes: La Reunion, Maurice, Rodriquez, Vol. 153–160. Sugar Industry Research Institute, Mauritius, 156 pp.

Radcliffe-Smith, A. (1986) Flora of Pakistan No. 176 Euphorbiaceae. University of Karachi, Karachi, Pakistan, 177 pp.

Rama Rao, M. (1914) Flora of Travancore. Government Press, Trivandrum, India, 448 pp.

Rao, R.S. & Sreeramulu, S. H. (1986) Flora of Srikakulam District. Hon. Secretary, Indian Botanical Society, Meerut, 640 pp.

Ray, J. (1688) Historiae Plantarum, Vol. 2. Henricum Faithorn, London, 1940 pp.

Redoute, P.J. (1799) Plantes Grasses, p. 46, plate 46. Garnerie Libraire, Paris, 236 pp.

Reddy, C.S., Reddy, K.N., Murthy, E.N., & Raju, V.S. (2009) Tree Wealth of Eastern Ghats of Andhra Pradesh, India: an updated checklist. Checklist 5 (2): 173–194.

Rheede tot Draakestein, H.A. van (1679) Hortus Malabaricus, Vol. 2. Johannis van Someren and Johannis van Dyck, Amsterdam, 110 pp.

Ridley, H.N. (1924) The Flora of the Malay Peninsula, Vol. 3. L. Reeve and Co., Ltd. London, 405 pp.

Roxburgh, W. (1812) Hortus Bengalensis. Mission Press, Serampore, India, 300 pp.

Roxburgh, W. (1832). Flora Indica, Vol. 2. W. Thacker and Co., Serampore, India, 587 pp.

Rumphius, G.E. (1743) Herbarium Amboinense, Vol. 4. Franciscum Changdion, Joannem Catuffe, Hermannum Uytwerf, Amsterdam, 154 pp.

Saldanha, C. J. & Nicolson, D.H. (1976) Flora of Hasan District, Karnataka, India. Amerind Publishing Co. Pvt. Ltd., New Delhi, 915 pp.

Saldanha, C. J. & Nicolson, D.H. (1996) Flora of Karnataka, Vol. 2. Oxford and IBH Publishing Co., New Delhi, 316 pp.

Sankara Rao, K., Singh, A.R., Kumar, D., Swamy, R.K., Page, N. (2016) Digital Flora of Eastern Ghats. Available from: http://floraeasternghats.ces.iisc.ac.in/herbsheet.php?id=1385&cat=4 (accessed 8 May 2020).

Sankara Rao, K., Swamy, R.K., Kumar, D., Singh, A.R., Bhat, K.G. (2019) Flora of Peninsular India. Available from: http://flora-peninsula-indica.ces.iisc.ac.in/herbsheet.php?id=3821&cat=7 (accessed 3 May 2020).

Sarojinidevi, N. (2017) *Euphorbia venkatarajui sp. nov.* (Euphorbiaceae) from Eastern Ghats of Andhra Pradesh, India. Nordic Journal of Botany. 35(3): 359–364.

Sarojinidevi, N. & Kullayiswamy, K.R. (2018) *Euphorbia belagaviensis sp. nov.* (Euphorbiaceae), from Karnataka state, India. Euphorbia World 14(1): 24–29.

Savage, S. (1945) A Catalogue of the Linnaean Herbarium. The Linnaean Society of London, London, 225 pp.

Saxena, H.O., Brahmam, M. (1995) The Flora of Orissa, Vol. 3. Orissa Forest Development Corporation Ltd., Bhubaneswar, 2008 pp.

Schmelzer, G.H. (2008) *Elaeophorbia drupifera*. *In*: Schmelzer, G.H. & Gurib-Fakim, A. (Eds.), Plant Resources of Tropical Africa (11(1), Medicinal Plants 1. PROTA Foundation, Wageningen, Netherlands, pp. 238 – 241.

Sealy, J.R. (1956) The Roxburgh Flora India Drawings at Kew. Kew Bulletin 11: 297– 348.

Seba, A. (1734) Locupletessimi Rerum Naturalum Thesauri, Vol. 1, Janssonia-Waesbergios, J. Wetstenium, & Gul. Smith, Amsterdam, 178 pp.

The Linnaean Collections (ND), Available from: https://linnean-online.org/4595/#?s=0&cv=0 (accessed 24 November 2022).

Singh, M. (2002). Succulent Plants of India, an introduction. Mrs. Meena Singh, Noida, 65 pp.

Standley, P.C. & Steyermark, J.A. (1949). Fieldiana: Botany, Flora of Guatemala, Vol. 24 (6). Chicago Natural History Museum, Chicago, 440 pp,

Staph, O. (1909) *Elaeophorbia drupifera* Staph. *In*: Prain, D. (Ed.), Hooker’s Icones Plantarum, Vol. 29. Dulau and Co., London, pp. 2801–2900.

Steudel, A.F. (1840) Nomenclator Botanicus seu: Synonymia Plantarum Universalis. J.G. Cottae, Stutgart, 810 pp.

Stewart, J.L. (1874) The Forest Flora of North-west and Central India. William H. Allen and Co., London, 608 pp.

Swamy, A.N. & Prasad, K. (2022) *Euphorbia ravii* (Euphorbiaceae: Subg. *Euphorbia*): a new species from Andhra Pradesh, India. Taiwania 67: 229–234.

Talbot, W.A. (1902) The Trees, Shrubs, and Woody Climbers of the Bombay Presidency. The Government Central Press, Bombay, 385 pp,

Thonning, P. (1827) *Euphorbia drupifera*. *In*: Schumacher, F.C. (Ed.), Beskrivelse af Guineiske Planter. Imprint of the Royal Danish Academy of Sciences and Letters, Copenhagen, 466 pp.

Tilli, M. (1723) Catalogus Plantarum Horti Pisani. Regiae Celcitudinis, Florence, 187 pp.

Turland, N.J., Wiersema, J.H., Barrie, F.R., Greuter, W., Hawksworth, D.L., Herendeen, P.S., Knapp, S., Kusber, W.-H., Li, D.-Z., Marhold, K., May, T.W., McNeill, J., Monro, A.M., Prado, J., Price, M.J. & Smith, G.F. (Eds.) 2018: International Code of Nomenclature for algae, fungi, and plants (Shenzhen Code) adopted by the Nineteenth International Botanical Congress Shenzhen, China, July 2017, p. 22. Regnum Vegetabile 159. Glashütten: Koeltz Botanical Books. DOI 10.12705/Code.2018

Van Royen, A. (1740) Flora Leydensis Prodromus. Samuelem Luchtmans, Leiden, 538 pp.

Warburg, O. (1894) Plantae Hellwigianae. *In:* Engler, A. (Ed.), Botanische Jahrbücher für Systematik, Planzengeschichte, und Plantsengeographie, Vol. 18, pp. 184–212.

Wight, (1852) Icones Indiae Orientales, Vol. 5 (2), p.19, plate 1862. P.R. Hunt, American Mission Press, Madras, India, pp. 1622–1920.

Wijnands, D. O. (1983) The Botany of the Commelins, A. A. Balkema, Rotterdam, 232 pp.

Wijnands, D. O. (1987) The Hortus Medicus Amstelodamensis. The Kew Magazine 4(2): 78–91.

Willdenow, C. L. (1799) Species Plantarum, Vol. 2(2). G. C. Nauk, Berlin, pp. 835–1340.

Wiman, J. (1752) No. 35, *Euphorbia*. In: C. Linnaeus, Amoenitates Academicae seu Dissertations etc. Vol. 111. Laurentii Salvii, Stockholm, pp. 100–131.

